# Adaptability and evolution of the cell polarization machinery in budding yeast

**DOI:** 10.1101/2020.09.09.290510

**Authors:** Fridtjof Brauns, Leila M. Iñigo de la Cruz, Werner K.-G. Daalman, Ilse de Bruin, Jacob Halatek, Liedewij Laan, Erwin Frey

## Abstract

How can a self-organized cellular function evolve, adapt to perturbations, and acquire new sub-functions? To make progress in answering these basic questions of evolutionary cell biology, we analyze, as a concrete example, the cell polarity machinery of *Saccharomyces cerevisiae*. This cellular module exhibits an intriguing resilience: it remains operational under genetic perturbations and recovers quickly and reproducibly from the deletion of one of its key components. Using a combination of modeling, conceptual theory, and experiments, we show that multiple, redundant self-organization mechanisms coexist within the protein network underlying cell polarization and are responsible for the module’s resilience and adaptability. Based on our mechanistic understanding of polarity establishment, we hypothesize how scaffold proteins, by introducing new connections in the existing network, can increase the redundancy of mechanisms and thus increase the evolvability of other network components. Moreover, our work suggests how a complex, redundant cellular module could have evolved from a more rudimental ancestral form.

## Introduction

Evolution is driven by an interplay of genotype mutations and selection operating on the level of biological function, that is, the phenotype. A mechanistic understanding of evolution therefore requires frameworks that connect the genotype to the phenotype (Rainey et al., 2017). When the phenotype (function) is determined by a self-organized process, the genotype-to-phenotype relation is not a simple one-to-one mapping (or “blueprint”). As a concrete example take intracellular (protein-based) pattern formation, which is essential for many essential cellular functions, like division and motility (Howard et al., 2011; Bi and Park, 2012; Chiou et al., 2017; Halatek et al., 2018; Ramm et al., 2019). The genotype determines the components (proteins), their interaction network and their copy numbers. Cellular function (the phenotype), on the other hand, emerges by the collective interplay of these components—governed by physical and chemical processes (diffusion, mass-action law) in the spatially extended cellular domain. A mechanistic understanding of the evolution of such collective (selforganized) functions has remained elusive so far (Johnson and Lam, 2010).

Here, we provide a concrete (and, to the best of our knowledge, first) example of how such understanding can be gained, using the cell Cdc42-polarization machinery of *Saccharomyces cerevisiae* (bud-ding yeast) as a model system. Cell polarization directs cell division of budding yeast through the formation of a polar zone with high Cdc42 concentration on the membrane (see Figure 1A-C). It is organized by a complex interaction network (Figure 1D) around the central polarity protein Cdc42, a GTPase that cycles between an active (GTP-bound) and an inactive (GDP-bound) state. The key features of these two states are that active Cdc42 is strongly membrane bound and recruits many down-stream factors, while inactive Cdc42-GDP can detach from the membrane to the cytosol where it diffuses freely.

**Figure 1.**
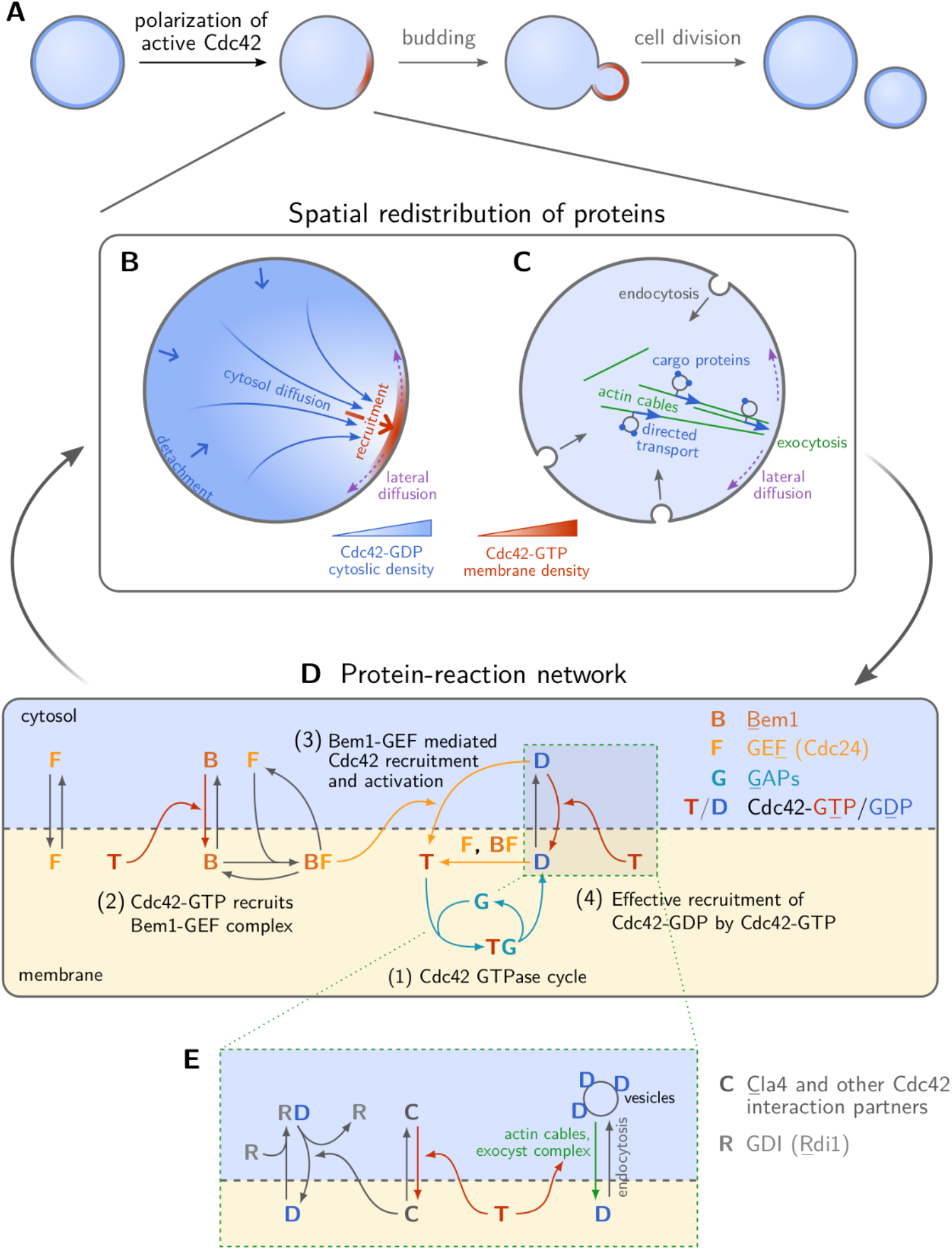
Cell division of *S. cerevisiae* is spatially controlled by self-organized polarization of Cdc42. **A** Starting from an initially homogenous distribution of Cdc42, a polar zone forms, marked by a high concentration of active Cdc42 on the plasma membrane. There are two pathways of directed transport in the cells: **B** Cytosolic diffusion becomes directed by spatially separated attachment (red arrow) and detachment (blue arrow) zones; **C** Vesicle transport (endocytic recycling) is directed along polar-oriented actin cables. Active Cdc42 directs both cytosolic diffusion (by recruiting downstream effectors that in turn recruit Cdc42) as well as vesicle transport (by recruiting Bni1 which initiates actin polymerization). **D** Molecular interaction network around the GTPase Cdc42, involving activity regulators (GEF, GAPs), and the scaffold protein Bem1. An effective recruitment term accounts for Cdc42-recruitment to the membrane directed by Cdc42-GTP facilitated by Cdc42-downstream effectors (**E**). Details of the model and the mathematical implementation are described in the SI Sections 1 and 2.

In wild-type (WT) cells, polarization is directed by upstream cues like the former bud-scar (Kang, 2001; Marston et al., 2001; Kozminski et al., 2003; Bi and Park, 2012). Importantly however, Cdc42 can polarize *spontaneously* in a random direction in the absence of such cues (Irazoqui et al., 2003; Wedlich-Soldner, 2003; Goryachev and Pokhilko, 2008). What are the elementary processes underlying spontaneous Cdc42 polarization? On the timescale of polarity establishment, the total copy number of Cdc42 proteins (as well as its interaction partners) is nearly constant. Hence, to establish a spatial pattern in the protein concentration, the so-called polar zone, the proteins need to be spatially redistributed in the cell by *directed transport*. There are two distinct, mostly independent, pathways for directed transport that have been established by experimental and theoretical studies (Wedlich-Soldner, 2003; Goryachev and Pokhilko, 2008; Freisinger et al., 2013; Klünder et al., 2013; Woods et al., 2015): cytosolic diffusion and vesicle-based active transport along polarized actin cables (Figure 1B,C). Once a polar zone has been established, the ensuing concentration gradient on the membrane leads to a diffusive flux of proteins away from the polar zone. To maintain the polar zone, this flux on the membrane must be counteracted continually by (re-)cycling the proteins back to the polar zone via a flux from the cytosol to the membrane (Goryachev and Pokhilko, 2008; Klünder et al., 2013; Chiou et al., 2017). In WT cells, Cdc42-GTP recruits Bem1 from the cytosol which in turn recruits Cdc24 (see Figure 1D) (Bose et al., 2001; Irazoqui et al., 2003). The membrane-bound Bem1-Cdc24 complex then recruits more Cdc42-GDP from the cytosol and activates it (nucleotide exchange) (Butty, 2002). The hallmark and crucial element of this *mutual recruitment mechanism* is the co-localization of Cdc42 and its GEF (Butty, 2002; Goryachev and Pokhilko, 2008; Howell et al., 2009; Kozubowski et al., 2008; Woods et al., 2015).

Deletion of Bem1 severely impedes the cells’ ability to polarize and bud (Chenevert et al., 1992; Irazoqui et al., 2003) by disrupting localized Cdc42 activation (Kozubowski et al., 2008; Woods et al., 2015). Intriguingly, in experimental evolution, *bem1*Δ mutants are reproducibly rescued by the subsequent loss of Bem3 (Laan et al., 2015). Bem3 is one of four known Cdc42-GAPs that catalyze the GTP-hydrolysis and hence switch Cdc42 into its inactive, GDP-bound state. The loss of Bem3 clearly does not replace Bem1 as it does not provide a scaffold between Cdc42 and Cdc24. How then does the loss of Bem3 rescue the pattern-forming capability of the *remaining* Cdc42-polarization machinery? Interestingly, it has been reported that *bem1*Δ cells can be rescued by fragments of Bem1 that do not interact with Cdc42-GTP but still bind to the membrane and to Cdc24 (Smith et al., 2013; Grinhagens et al., 2020). These Bem1 fragments therefore cannot mediate mutual recruitment of Cdc42 and its GEF Cdc24, but only confer increased global (homogeneous) GEF activity by relieving Cdc24’s autoinhibition (Shimada et al., 2004; Rapali et al., 2017). This suggests that Cdc42 polarization can emerge *independently* of GEF co-localization, but the underlying mechanism remains unclear.

The adaptability of budding yeast’s cell polarization module makes it an ideal model system for studying the evolution of self-organized function. Here, we develop a theory that shows that this cellular module comprises multiple redundant reaction–diffusion mechanisms. It reveals that in addition to the Bem1-mediated mutual recruitment mechanism, a distinct and latent mechanism exists in the Cdc42-polarization machinery. We show that this latent mechanism operates under different constraints on the protein copy numbers than the wild-type mechanism and is activated by the loss of Bem3 which lowers the total copy number of GAPs. This explains how cell polarization is rescued in *bem1*Δ *bem3*Δ cells (Laan et al., 2015), and also reconciles the puzzling experimental findings outlined above. Moreover, we experimentally confirm the predictions of our theory on how cell polarization in various mutants can be rescued by changing the Cdc42 copy number. On the basis of the mechanistic understanding of the cell polarization module in budding yeast, we then propose a possible evolutionary scenario for the emergence of this self-organized cellular function. We formulate a concrete hypothesis how evolution might leverage scaffold proteins to introduce new connections in an existing network, and thus increase redundancy of mechanisms within a functional module. This redundancy loosens the constraints on the module and thereby enables further evolution of its components, for instance by duplication and sub-functionalization (Magadum et al., 2013).

## Results

As basis for our theoretical analysis, we first need to formulate a mathematical model of the cells’ Cdc42-polarization machinery that is able to explain Bem1-independent polarization. The interplay of spatial transport processes (Figure 1B,C) and protein-protein interactions (Figure 1D) is described in the framework of reaction–diffusion dynamics. The biochemical interaction network we propose is based on the quantitative model introduced in (Klünder et al., 2013) and extends it in several important ways. It accounts for the Cdc42 GTPase cycle and the interactions between Cdc42, Bem1 and Cdc24 (Goryachev and Pokhilko, 2008). Importantly — extending previous models — we explicitly incorporate the transient formation of a GAP-Cdc42 complex as an intermediate step in the enzymatic interaction between GAPs and Cdc42 (Zhang et al., 1997). In addition, we include effective self-recruitment of Cdc42-GDP to the membrane which is facilitated by membrane-bound Cdc42-GTP. This effective recruitment accounts for vesicle-based Cdc42 transport along actin cables (Slaughter et al., 2009; Layton et al., 2011; Freisinger et al., 2013) and putative recruitment pathways mediated by Cdc42GTP downstream effectors such as Cla4 and Gic1/2 (Tiedje et al., 2008; Das et al., 2012; Daniels et al., 2018). A detailed description of the model, illustrated in Figure 1D, and an in-depth biological motivation for the underlying assumptions are given in the SI Section 1.

### The Cdc42 interaction network facilitates a latent polarization-mechanism

We first ask whether the proposed reaction–diffusion model of the Cdc42 polarization machinery can explain spontaneous polarization in the absence of Bem1, i.e. without GEF co-localization with Cdc42. To this end, we perform a linear stability analysis of the model which identifies the regimes of selforganized pattern formation. A large-scale parameter study (see SI Section 5) reveals that in the absence of Bem1 there is a range of protein numbers of Cdc42 and GAP where polar patterns are possible (Figure 2B), i.e. that there is a latent polarization mechanism. However, in contrast to the Bem1-dependent mutual recruitment mechanism (Figure 2A), we find that the regime of operation for this latent mechanism is more limited and requires a sufficiently low GAP/Cdc42-concentration ratio (Figure 2B).

**Figure 2.**
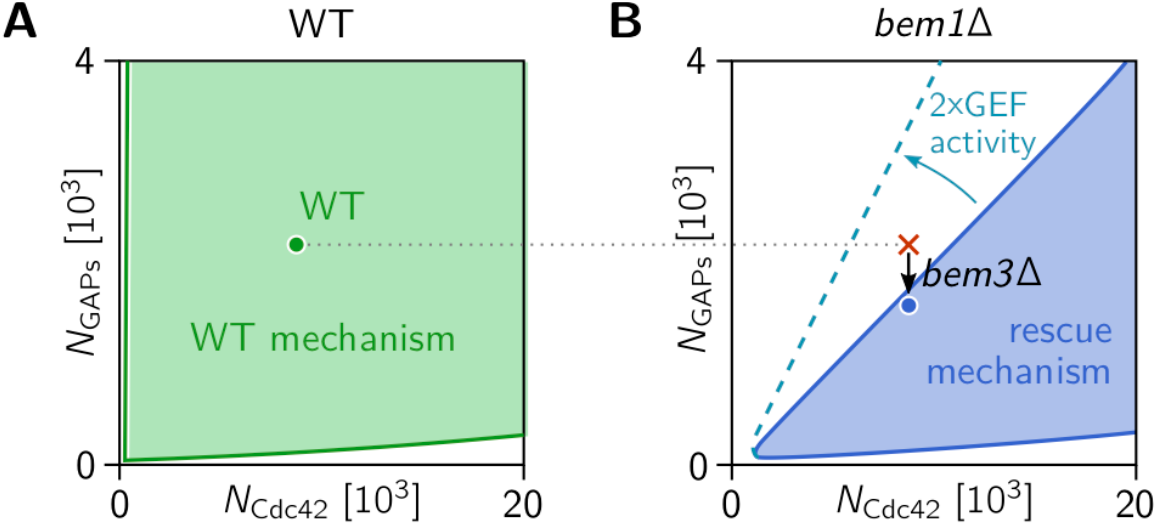
Regimes of operation of the Bem1-mediated wild-type mechanism and the latent mechanism for cell polarity. Stability diagrams as a function of GAP- and Cdc42 concentrations in presence and absence of Bem1 obtained by linear stability analysis (see SI Section 3) of the mathematical model for the Cdc42-polarization machinery (see Figure 1 and SI Section 2). Shaded areas indicate regimes of lateral instability, i.e. where a spontaneous polarization is possible. **A** In WT cells, the scaffold protein Bem1 is present and facilitates spontaneous polarization by a mutual recruitment mechanism that is operational in a large range of Cdc42 and GAP concentrations (Goryachev and Pokhilko, 2008; Klünder et al., 2013). The green point marks the Cdc42 and GAP concentrations of WT cells. **B** In the absence of Bem1, spontaneous polarization is restricted to a much smaller parameter-space region in our model, because the regime of operation of the Bem1-indepenendent mechanism is inherently is delimited by a critical ratio of GAP concentration to Cdc42 concentration. The Cdc42 and GAP concentrations of *bem1*Δ cells and *bem1*Δ *bem3*Δ are marked by the red cross and blue point, respectively. The experimental observation that *bem1*Δ cells do not polarize, whereas *bem1*Δ *bem3*Δ polarize can be used to infer a range for the critical GAP/Cdc42-concentration ratio. Increasing the GEF activity of Cdc24 increases this critical ratio (dashed blue line). (The model parameters were obtained by sampling for parameter sets that are consistent with the experimental findings on various mutants, as described in detail in SI Section 5.)

What is the mechanistic cause for this constraint? To answer this question, we need to understand how the Cdc42-polarization mechanism works in the absence of Bem1. As emphasized above, Cdc42-polarization requires two essential features—directed transport of Cdc42 to the polar zone and localized activation of Cdc42 there. The first feature, directed transport, is accounted for in the model by effective recruitment of Cdc42-GDP to the membrane mediated by active Cdc42 (Figure 1D).

### GAP saturation can localize Cdc42 activity to the polar zone

How is the second feature, localization of Cdc42 activity to the polar zone, implemented in the absence of Bem1? Instead of directly increasing the rate of Cdc42 activation in the polar zone (via recruitment of the GEF Cdc24 by Bem1), localization of activity can also be achieved by decreasing the rate of Cdc42 *deactivation* in the polar zone and increasing it away from the polar zone. In fact, if enzyme saturation limits the net deactivation rate, a simple increase in Cdc42 density *generically* leads to a decrease of the Cdc42 deactivation rate (per Cdc42 molecule). Enzyme saturation of catalytic reactions occurs when the dissociation of the transient enzyme-substrate complex (here the GAP-Cdc42 complex) is the rate limiting step such that the enzymes are transiently sequestered in enzyme-substrate complexes. Indeed, it has been shown that this is the case for GAP-catalyzed hydrolysis of Cdc42 in budding yeast (Zhang et al., 1997). Furthermore, enzyme saturation requires that a large fraction of enzymes is sequestered in enzyme–substrate complexes, i.e. that the total enzyme density is sufficiently low compared to the substrate density, as we found in the linear stability analysis (Figure 2B).

In summary, GAP saturation localizes Cdc42 activity to the polar zone, by decreasing the deactivation rate in the polar zone, where Cdc42 density is high, relative to the remainder of the membrane, where Cdc42 density is low. This, in conjunction with transport of Cdc42 to the polar zone, drives spontaneous cell polarization. Interestingly, enzyme saturation of Cdc42 hydrolysis is one of the six theoretically possible mechanisms for pattern formation that were hypothesized by a generic mathematical analysis of feedback loops in GTPase cycles (Goryachev and Leda, 2017).

### The latent polarization-mechanism explains the rescue of Bem1 deletes

The Bem1-independent rescue mechanism requires a sufficiently low GAP/Cdc42-concentration ratio to be functional (Figure 2B). This suggests that *bem1*Δ cells are not able to polarize because their GAP copy number is too high. Our model predicts that the loss of GAPs can rescue cell polarization by bringing their total copy number into a regime where the Bem1-independent mechanism is operational, as indicated by the arrow in Figure 2B. This is in accordance with evolution experiments showing that *bem1*Δ cells are reproducibly rescued by a subsequent loss-of-function mutation of the GAP Bem3 (Laan et al., 2015). Bem3 accounts for approximately 25% of the total copy number of all Cdc42-GAPs (Kulak et al., 2014), indicating that *bem1*Δ mutants are close to the GAP/Cdc42-ratio threshold of the Bem1-independent mechanism. This proximity of the protein copy numbers to the threshold explains why a low fraction (about 1 in 10^5^) of mutants are able to polarize and divide, after *BEM1* has been deleted (Laan et al., 2015): Protein expression levels vary stochastically from cell to cell such that a small fraction of cells lies in the concentration regime where the latent polarization mechanism drives spontaneous cell polarization.^1^

Rather than by the loss of a GAP, the GAP/Cdc42-concentration ratio could also be brought down by an increase of the Cdc42 copy number. Yet another option would be an increase of Cdc24’s GEF activity which would increase the critical threshold in GAP/Cdc42-concentration ratio (see dashed line in Figure 2B). However, compared to a loss-of-function mutation, such mutations have a much smaller mutational target size and are therefore much less frequent. Moreover, one might wonder why it is specifically Bem3, rather than one of the other GAPs, that is lost to rescue the *bem1*Δ strain. Some hints to answer this outstanding question are provided by a detailed theoretical analysis of the rescue mechanism discussed below (***Functional submodules of cell polarization***).

### Experiments confirm theoretical predictions

Based on the GAP/Cdc42-ratio constraint in the rescue mechanism, our theory makes two specific predictions: (i) Increasing the copy number (i.e. overexpression) of Cdc42 will rescue cell polarization of *bem1*Δ cells by invoking the Bem1-independent mechanism. (ii) Polarization of *bem1Δbem3Δ* cells will break down if the expression level of Cdc42 is lowered compared to the WT level (Figure 2B).

To test these model predictions experimentally, we first constructed different yeast strains with Cdc42 under an inducible galactose promoter such that we can tune the Cdc42 copy number by varying the galactose concentration in the growth media (Yocum et al., 1984): a *bem1Δ* strain (yWKD069), a *bem1Δ bem3Δ* (yWKD070), and a modified WT strain (yWKD065) (see ***Materials and Methods***).

As a next step, we inoculated the different strains at varying galactose concentration in 96 well plates, that were placed in a plate reader to measure the cell density over time, and thereby determined the growth rate (see ***Materials and Methods***). For every galactose concentration, the growth rates are normalized to those of WT cells, with Cdc42 under its native promotor (yLL3a), grown at the same galactose concentration. In Figure 3A the normalized growth rates of the different mutants are plotted. As expected, WT cells grow at all galactose concentrations. In contrast, WT cells with Cdc42 under the galactose promotor (yWKD065), do not grow in the absence of Cdc42 (0% galactose concentration), since a failure to polarize severely impairs cell division and eventually leads to cell death and thus zero growth rate (Irazoqui et al., 2003). Our data show that the WT mechanism is rather insensitive to Cdc42 copy number, even for very low expression of Cdc42, in accordance with theory (Figure 2A).

**Figure 3.**
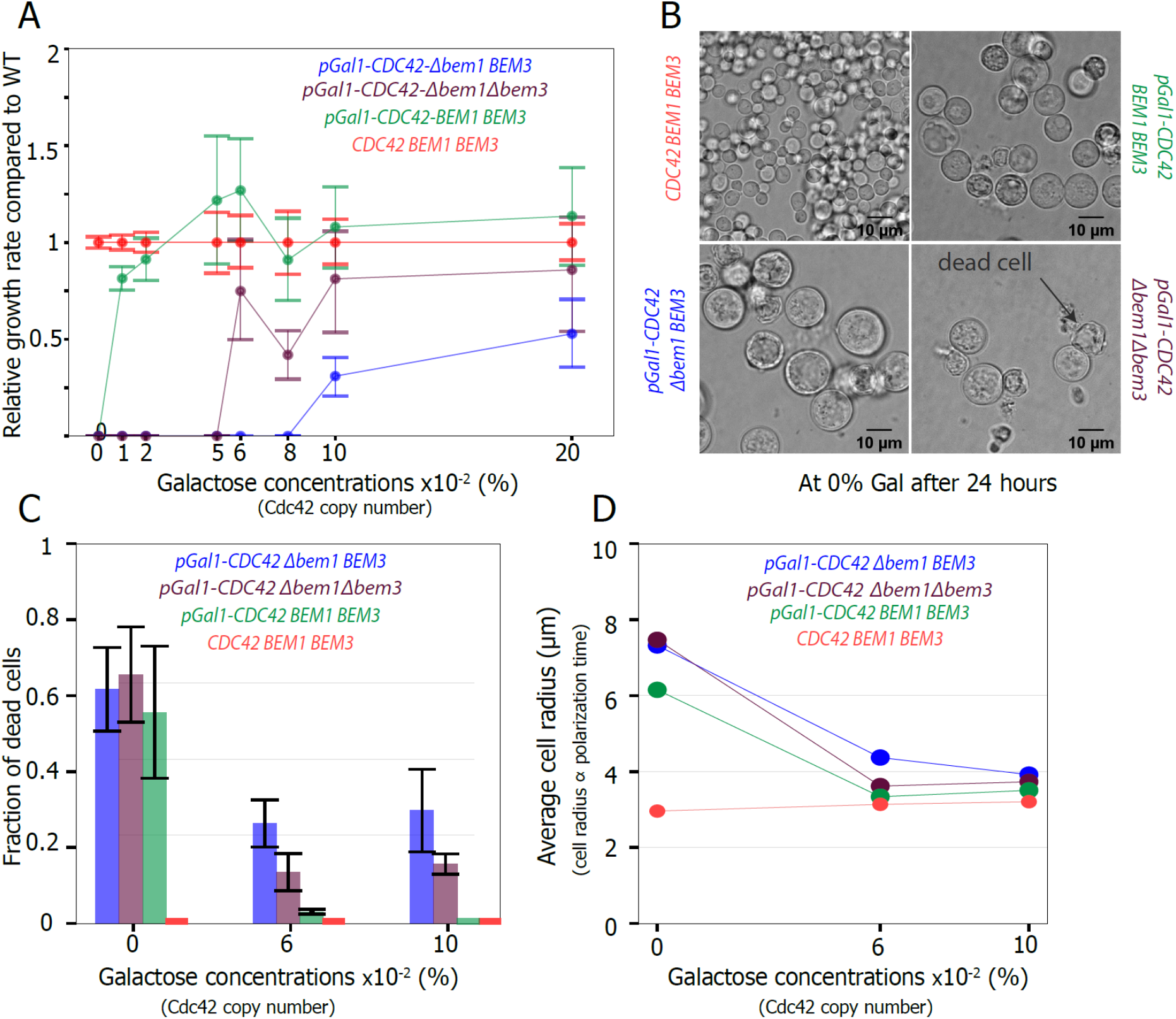
Experiments confirm theoretically predicted effect of Cdc42 copy number on the latent polarity-mechanism. **A** Growth rate of the different mutants (relative to the growth rate of WT cells with Cdc42 under its native promotor at that galactose concentration (red)) against the galactose concentration galactose concentration (proxy for Cdc42 copy number) show that higher expression of Cdc42 rescues *bem1*Δ cells and to a lesser extent *bem1*Δ *bem3*Δ cells; the error bar indicates the 68% credible interval, see materials and methods). The spread in the data is partially caused by experimental errors, as well by demographic noise, i.e. new fitter mutants arising by random mutations and taking over the population. **B** Microscopy images shows the morphology of dead and alive cells for the different strains after 24 hours at 0% galactose concentration, resulting in a Cdc42 copy number that approximates zero; scale bar indicates 10 μm. **C** The fraction of dead cells for different mutant strains vs galactose concentration shows that increasing Cdc42 copy number reduces cell death in *bem1*Δ and *bem1*Δ *bem3*Δ cells; the error bar indicates the standard error of the mean. **D** The average cell radius (proxy for polarization time) versus the galactose concentration shows that increasing Cdc42 copy number reduces polarization time; the error bar, which is the standard error of the mean, is smaller than the data symbol.

Our model predicts that *bem1Δ* cells need the highest Cdc42 copy number to polarize, WT cells will need the least, and the *bem1Δ bem3Δ* cells should be in between. We indeed find that the *bem1Δ* strain (yWKD069) grows in media with 0.1% or higher galactose concentration. We did inoculate these strains at lower galactose concentration, but never observed any growth (*n* ≥ 2 experiments, with 4 technical replicates per condition). The *bem1Δbem3Δ* cells (yWKD070) grow only in a galactose concentration of at least 0.06%. For WT cells with Cdc42 under the galactose promotor growth we observe and reduced growth rate at 0.01% galactose concentration but growth is only fully inhibited at 0% galactose concentration. All of the above experimental observations agree with our specific theoretical predictions. To show that the differences in population growth rates are directly caused by the ability of cells to polarize, rather than for example pleiotropic changes in another cell cycle phase, we performed a second set of experiments, where we measured the cell radius using light microscopy (Figure 3B). It was previously shown that the cell radius correlates linearly with the time it takes for cells to polarize (Allard et al., 2018; Laan et al., 2015): cells that take longer to polarize are on average larger than cells that polarize fast because yeast cells continue to grow during polarity establishment, allowing us to use the cell radius as a proxy for the polarization time. Additionally, we verified that, at low Cdc42 copy numbers, cells cannot polarize at all and thus die. Consistent with the population growth data, we observed that after 24 hours at 0% galactose concentration, for every genetic background where Cdc42 is under the galactose promotor, the vast majority of cells are not able to polarize or polarize very slowly, because they are either dead (Figure 3B,C) or very large (Figure 3B,D). We also confirm that the average cell radius (and thus the polarization time) and death rate of cells with Cdc42 under its native promotor are not affected by the galactose concentration (Figure 3C,D in red). At 0.06% galactose concentration, *bem1*Δ *bem3*Δ the cells’ radii (and thus polarization times) are closer to WT cell radii than those of *bem1*Δ cells. This agrees with the population growth data. And at 0.1% Gal the average cell radius for live cells for all mutants were approximately equal to the average WT cell radius (Figure 3D). Interestingly, after 24 hours at 0% galactose concentration, WT cells with Cdc42 under the galactose promotor are still polarizing faster than the *bem1*Δ and the *bem1*Δ *bem3*Δ cells, as indicated by their smaller average cell radius (Figure 3D). This observation confirms our above observation that a very small number of Cdc42 molecules is sufficient for WT cells to polarize and thus for the WT mechanism to be operational.

Taken together, the experimental data confirm the theoretical prediction that the Bem1-independent rescue mechanism is operational only below a threshold GAP/Cdc42-concentration ratio. In addition, we find that the Bem1-dependent WT mechanism is surprisingly insensitive to Cdc42 copy number, i.e. operates also at very low Cdc42 concentration. This significant difference in Cdc42 copy number sensitivity between the WT mechanism and the rescue mechanism is in the context of our theory explained by the qualitative difference of their principles of operation, as we discussed above in the section ***The Cdc42 interaction network facilitates a latent polarization mechanism***. While the WT mechanism is based on recruitment of the GEF Cdc24 to the polar zone, mediated by the scaffold protein Bem1, the rescue mechanism crucially involves enzyme saturation of Cdc42 hydrolysis due to high Cdc42 density in the polar zone. This enzyme saturation requires a sufficiently large Cdc42 copy number relative to the GAP copy number. In the section ***Functional submodules of cell polarization*** below, we will analyze the mathematical model, and the qualitative and conceptual differences between these two mechanisms in more detail.

### The latent rescue mechanism explains and reconciles previous experimental findings

In previous experiments, several Bem1 mutants were studied that perturb Bem1’s ability to mediate co-localization of Cdc24 to Cdc42-GTP, the key feature that underlies operation of the WT mechanism (Howell et al., 2009; Smith et al., 2013; Bendezú et al., 2015; Woods et al., 2016; Witte et al., 2017; Grinhagens et al., 2020). The observations from these experiments have remained puzzling and apparently conflicting among one another as of yet. As we show in detail in the Supplementary Discussion in SI Section 6, the latent rescue mechanism predicted by our mathematical model explains and reconciles all of these previous experimental findings. The key insight is that the latent rescue mechanism can be activated by a global increase of GEF activity (see dashed line in Figure 2B). Bem1 mutants that lack the Cdc42-interaction domain but still bind to the GEF Cdc24 may provide such a global increase of GEF activity and thus rescue polarization of *bem1*Δ cells. Moreover, in accordance with optogenetics experiments (Witte et al., 2017), our mathematical model predicts that outside the regime of spontaneous polarization, the latent, Bem1-independent mechanism can also be induced by a sufficiently strong local perturbation of the membrane-bound GEF concentration.

### Functional submodules of cell polarization

Cell polarization in budding yeast is a functional module based on a complex protein interaction network with Cdc42 as the central polarity protein (cf. Figure 1B-D). As we discuss next, the full network can be dissected into *functional submodules*. Here, the term functional submodule refers to a *part* of the full interaction network with a well-defined function in one or more pattern-forming mechanisms. Our theoretical analysis will reveal that an interplay of two (or more) *functional submodules* each constitutes a fully functional cell polarization mechanism.

As we argued in the ***Introduction***, establishment and maintenance of cell polarity requires that Cdc42-activity is localized to membrane regions with a high density of Cdc42. This can be achieved in two different ways. First, by the recruitment of the scaffold protein Bem1 to Cdc42-GTP, which in turn recruits the GEF (Cdc24) and thus localizes Cdc42 activation to the polar zone, where Cdc42 density is high (Figure 4A, top left). We call this the *polar activation* submodule. Second, GAP saturation in regions of high local Cdc42 densities can localize Cdc42 activity to the polar zone (Figure 4A, top right), as described above in the subsection ***GAP saturation can localize Cdc42 to the polar zone***. The transient sequestration of GAPs in Cdc42-GAP complexes is essential for this *polar GAP saturation* submodule. The third submodule (Figure 4A, bottom) that we term *Cdc42 transport*, comprises various modes of Cdc42 transport towards the polar zone: vesicle transport along polarized actin cables (cf. Figure 1B) and effective (self-)recruitment of Cdc42 from the cytosol. Several experiments indicate that downstream effectors of active Cdc42, such as Cla4, Gic1 and Gic2 may provide such effective recruitment in the absence of Bem1 (Tiedje et al., 2008; Kang et al., 2018; Daniels et al., 2018).

**Figure 4:**
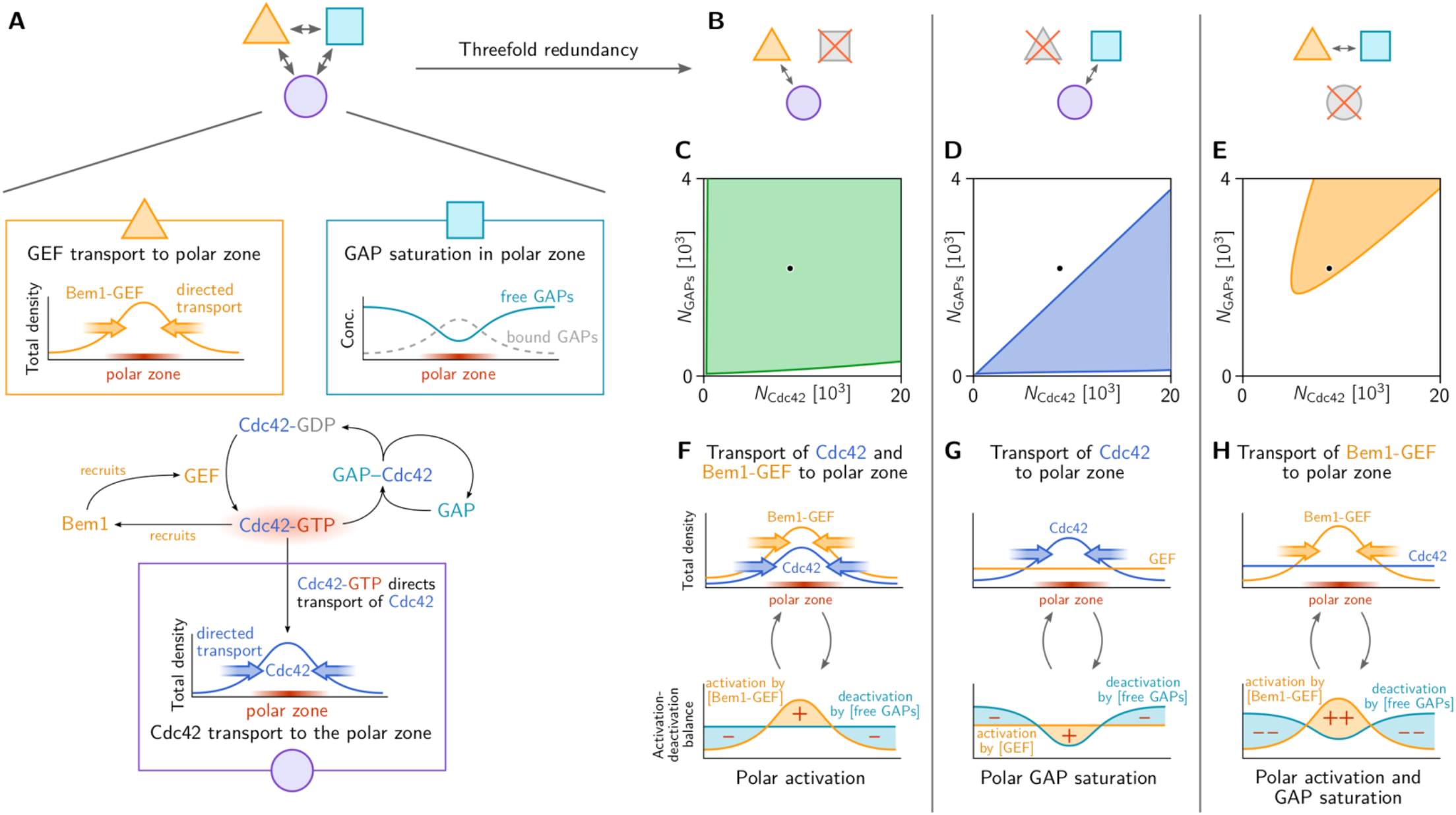
Three functional submodules constitute three distinct mechanisms of Cdc42-GTP polarization. **A** Three functional submodules of the Cdc42 interaction network contribute to the formation and maintenance of a *polar zone* (region of high Cdc42-GTP concentration, highlighted in red): (i) Transport of Cdc42 towards the polar zone. High Cdc42 activity can be maintained due to (ii) GAP saturation in the polar zone and by (iii) transport of the GEF to the polar zone via the scaffold protein Bem1. **B** Combinations of pairs of these functional submodules constitute mechanisms of self-organized pattern formation. **C** –**E** These mechanisms are operational in different regimes of the total copy number of Cdc42 and GAPs. The WT mechanism (**F**) is largely insensitive to copy number variations (**C**) because it based on mutual recruitment of Cdc42 and Bem1-GEF complexes, and does not depend on saturation of GAPs in the polar zone. In contrast, when the GEF is not transported to the polar zone (e.g. due to a deletion of Bem1), only GAP saturation in the polar zone maintains high Cdc42 activity there, while deactivation dominates away from the polar zone. Therefore, the polarization mechanism (**G**) is sensitive to the GAP copy number (**D**). **H** Remarkably, if transport of Cdc42 is suppressed, e.g. by strongly binding it to the membrane, a combination of Bem1-GEF complex recruitment and polar GAP saturation maintain a localized high Cdc42 activity even though the total density of Cdc42 is homogenously distributed.

**Figure 5.**
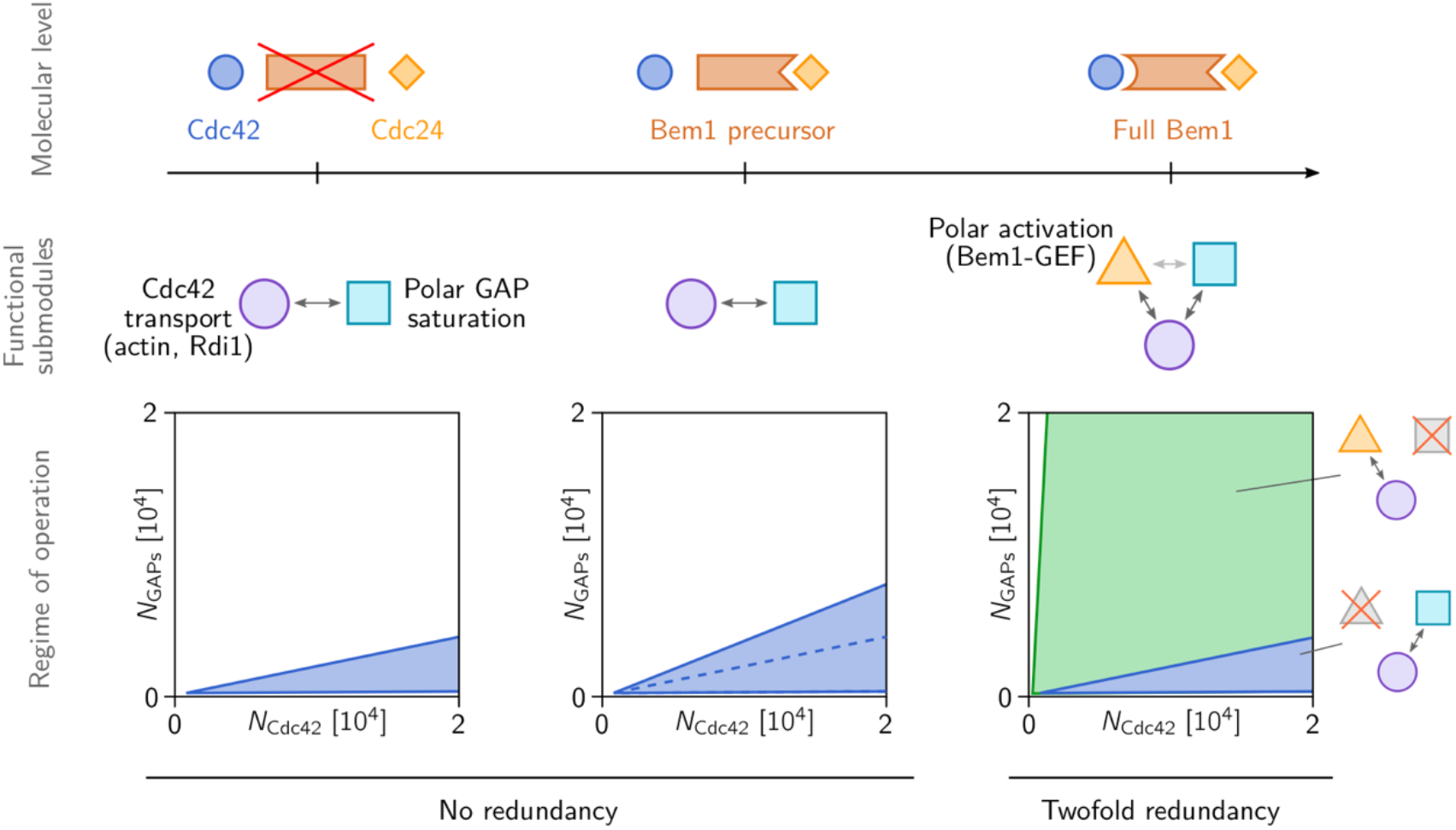
Hypothetical evolution of Bem1. (*Left*) The Bem1-independent “rescue” mechanism based on GAP saturation and Cdc42 transport towards membrane bound Cdc42-GTP is operational only in a limited range of the GAP/Cdc42-concentration ratios (cf. Figure 4D). (*Center*) a Bem1 precursor (Bem1-fragment) that binds to Cdc24 and relieves its auto-inhibition increases the range of viable GAP/Cdc42-concentration ratios and thus increases the robustness against copy number variations (cf. Figure 2). It does, however, not change the underlying mechanism qualitatively. (*Right*) Domain fusion of a Cdc42-GTP-binding domain with the Cdc24-binding Bem1-precursor, leads to a new connection in the Cdc42-interaction network that leads to recruitment of Cdc24 to the polar zone. On the level of sub-modules, this new connection constitutes a new functional submodule that we called “polar activation” (yellow triangle). In conjunction with transport of Cdc42 towards the polar zone, polar activation gives rise to the highly robust mutual-recruitment mechanism that is operational in WT yeast (regime of operation shaded in green in the (*N*_D_,*N*_G_)-parameter plane; cf. Figure 4C).

These three functional submodules represent different mechanistic aspects of the Cdc42-interaction network. Each submodule is operational only under specific constraints on the biochemical properties and copy numbers of the involved proteins. In the following, we exploit these constraints to study the roles of the submodules in the mathematical model by disabling them one at a time. This allows us to tease apart the mechanisms that are operational under the corresponding experimental conditions. The first submodule, *polar activation*, is disabled by the knock-out of Bem1. The second submodule, *polar GAP saturation*, is suppressed if the copy number of GAPs is too high. Alternatively, polar GAP saturation is rendered non-operational if the dissociation rate of the GAP-Cdc42 complex is too fast, or if the free GAPs diffuse very fast making additional free GAPs readily available in the polar zone. The third submodule, *Cdc42 transport*, can be switched off by immobilizing Cdc42, i.e. suppressing its spatial redistribution. Experimentally, this has been achieved in fission yeast by fusing Cdc42 to a transmembrane protein that strongly binds to the membrane and is nearly immobile there (Bendezú et al., 2015).

By performing linear stability analysis for the full mathematical model under each of these perturbations disabling one of the submodules at a time (as described in detail in SI Section 5, SI Table S4), we find that the remaining two submodules operate in concert to constitute a mechanism for spontaneous Cdc42 polarization, as illustrated in Figure 4B. Figures 4C-E shows the regime of operation of the three different mechanisms as a function of the total Cdc42 and GAP concentrations. Figure 4F-H illustrate the concerted interplay of directed protein-transport and regulation of Cdc42 activity (activation/deactivation) that underlie Cdc42-polarization in these three mechanisms.

Before we turn to the detailed descriptions of these mechanisms, we note that if two submodules are disabled simultaneously, the remaining submodule alone cannot facilitate pattern formation. In particular, and perhaps somewhat counterintuitively, self-recruitment of Cdc42 alone is not sufficient to drive spontaneous cell polarization (Altschuler et al., 2008; Goryachev and Leda, 2017).

### Wild-type mechanism: *Cdc42 transport plus polar activation*

The interplay of the Cdc42 transport submodule and the Cdc42-Bem1-Cdc24 recruitment submodule (polar activation), illustrated in Figure 3F, constitutes the WT mechanism that operates via mutual recruitment of Cdc42 and Bem1 (Irazoqui et al., 2003; Klünder et al., 2013; Freisinger et al., 2013). Characteristic for this mechanism is the co-localization of Cdc24 and Cdc42-GTP in the polar zone, as observed in previous experiments (Woods et al., 2016; Witte et al., 2017). Other than the rescue mechanism, the mutual recruitment mechanism does not require polar GAP saturation. Therefore, it is insensitive against high concentration of GAPs, i.e. it is operational for much higher GAP/Cdc42-concentration ratios than the rescue mechanism. Furthermore, it is robust against high diffusivity of free GAPs and high catalytic rates of the GAPs (fast decay of GAP-Cdc42 complexes into free GAP and Cdc42-GDP). This implies that in mathematical models of the WT mechanism the GAPs can be accounted for *implicitly* by a constant and homogeneous hydrolysis rate, as e.g. in (Goryachev and Pokhilko, 2008; Klünder et al., 2013; Kuo et al., 2014; Woods et al., 2016).

### Rescue mechanism: *Cdc42 transport plus polar GAP saturation*

The interplay of Cdc42 transport (including effective self-recruitment via actin and/or other down-stream effectors like Cla4) and GAP saturation in the polar zone, illustrated in Figure 3G, constitutes the latent, Bem1-independent rescue mechanism. Characteristic for this mechanism is that it does not require co-localization of Cdc24 to Cdc42-GTP in the polar zone (see Figure 4G). This lack of Cdc24 polarization would serve as a clear indicator of the rescue mechanism in future experiments using fluorescently labelled Cdc24. As explained above, the rescue mechanism relies on GAP saturation in the polar zone to maintain high Cdc42 activity there. This GAP saturation is suppressed by either high abundance, high catalytic activity, or fast transport (by cytosolic diffusion or vesicle recycling) of the GAPs.

The last constraint provides a plausible explanation why it is specifically Bem3 that needs to be deleted to rescue *bem1*Δ cells. In contrast to Rga1 and Rga2, Bem3 has been found to be highly mobile, probably because it cycles through the cytosol (Mukherjee et al., 2013). GAP saturation, i.e. the depletion of free GAPs in the polar zone, entails a gradient of the free GAP density towards the polar zone. A mobile GAP species like Bem3 will quickly diffuse along this gradient to replenish the free GAPs in the polar zone, relieving the GAP saturation there, and thus counteract the activation of Cdc42 in the incipient polar zone. Therefore, the loss of Bem3, rather than one of the other, less mobile GAPs, promotes the formation of a stable polar zone.

Interestingly, the formation of Min-protein patterns in *E. coli* relies on the same type of mechanism as the rescue mechanism for Cdc42-polarization: self-recruitment of an ATPase (MinD) and enzyme saturation of the AAP (MinE) that catalyzes MinD’s hydrolysis and subsequent membrane dissociation (Huang et al., 2003; Halatek and Frey, 2012; Halatek et al., 2018). The transient MinDE complexes play the analogous role to the Cdc42-GAP complexes here: In regions of high MinD density, MinE is sequestered in MinDE complexes, which limits the rate of hydrolysis until the complexes dissociate or additional MinE comes in by diffusion. Because MinE cycles through the cytosol, it rapidly diffuses into the polar zone where the density of free MinE is low, thus relieving the enzyme saturation there and eventually leading to a reversal of the MinD polarity direction. The repeated switching of MinD polarity due to redistribution of MinE is what gives rise to the Min oscillations in *E. coli*. Recently also stationary Min patterns have been observed *in vitro* (Glock et al., 2019). Conversely, oscillatory Cdc42 dynamics are found in the fission yeast *S. Pombe* (Das et al., 2012), and have also been indirectly observed in budding yeast mutants (Kuo et al., 2014; Ozbudak et al., 2005).

### Polarization with immobile Cdc42: Bem1-mediated recruitment plus polar GAP saturation

The interplay of Cdc42-Bem1-Cdc24 recruitment (polar activation) and the polar GAP saturation, illustrated in Figure 3H, facilitates polarization of Cdc42 activity without the spatial redistribution Cdc42’s total density. Instead, the proteins that are being redistributed are Bem1 and GEF. The polar zone is characterized by a high concentration of membrane-bound Bem1–GEF complexes which locally increase Cdc42 activity. Cdc42-GTP, in turn, recruits further Bem1 and GEF molecules to the polar zone. Characteristic for this mechanism is that Cdc42-GTP is polarized while the total Cdc42 density remains uniform on the membrane. Experimentally, this has been observed in fission yeast using Cdc42 fused to a transmembrane domain (Cdc42-psy1™) that renders Cdc42 nearly immobile. The polarization machinery of fission yeast is closely related to the one of budding yeast; it operates based on the same mutual recruitment pathway with Scd1 and Scd2 taking the roles of Cdc24 and Bem1 (Chiou et al., 2017). In future experiments, it would be interesting to test whether the Cdc42-psy1™ also facilitates polarization in budding yeast (potentially in a strain with modified GAP or Cdc42 copy number as the regime of operation might not coincide with the WT copy numbers).

## Conclusion and discussion

“How do cells work and how did they come to be the way they are?” (Lynch et al., 2014) We have approached this fundamental question of evolutionary cell biology by analyzing in depth a concrete system — the Cdc42 polarization machinery of budding yeast — that plays an essential role in the cell division of this model organism. Previous experiments showed that this machinery exhibits an intriguing resilience. It remains operational under many experimental (genetic) perturbations (Brown et al., 1997; Smith et al., 2013; Woods et al., 2015; Bendezú et al., 2015; Witte et al., 2017; Grinhagens et al., 2020), and recovers quickly and reproducibly from the deletion of one of its key components, the scaffold protein Bem1 (Laan et al., 2015). In the following we will shortly recapitulate the main insights we gained by studying the cell biology of this system and then show how a mechanistic understanding of self-organized cellular function can lead to fundamental insights into the way this function could have evolved from a more rudimental ancestral form.

### Mechanistic understanding of the cell polarization module in budding yeast

We have discovered that multiple, redundant self-organization mechanisms coexist within the protein network underlying cell polarization, that are responsible for the resilience and adaptability of the cell polarization module. By dissecting the full cellular polarization module into *functional submodules* we have identified distinct mechanisms of self-organized pattern formation, including the wild-type mechanism relying on the colocalization of Cdc42 with its GEF and a latent Bem1-independent rescue mechanism. Our theory, confirmed by experimental analysis, reveals that these mechanisms share many components and interaction pathways of this network. This implies that the redundancy of cell polarization is not at the level of individual components or interactions but arises on the level of the emergent function itself: If one submodule is rendered non-functional, the combination of the remaining submodules still constitutes an operational mechanism of cell polarization — if parameters, in particular protein copy numbers, are tuned to a parameter regime where these remaining submodules are operational. Redundancy hence provides adaptability — the ability to maintain function despite (genetic) perturbations, like the knockout of Bem1.

### The physics of self-organization imposes constraints on evolution

Our theoretical and experimental results highlight the importance of protein copy numbers as control parameters that decide whether a mechanism of spontaneous cell polarization is operational. Phrased from a genetic perspective, the genes that code for components of the cell polarization machinery are *dosage sensitive* (Papp et al., 2003). On the one hand, this entails that mutations of cis-regulatory elements (like promoters and enhancers) (Wittkopp and Kalay, 2012) can tune the copy numbers of proteins to the regime of operation of a specific cell-polarization mechanism and optimize the function within that regime. On the other hand, dosage sensitivity constrains evolution of the polarization-machinery’s components via duplication and sub-functionalization (Conant and Wolfe, 2008; Papp et al., 2003).

One of our key findings is that the constraints on a single particular mechanism can be circumvented by the coexistence of several redundant mechanisms of self-organization that operate within the same protein-interaction network. The regimes of operation — and, hence the dosage sensitivity of specific genes — can differ vastly between these distinct mechanisms. Therefore, redundancy on the level of mechanisms allows the module’s components to overcome constraints like dosage sensitivity and thus promotes “evolvability” — the potential of components to acquire new (sub-)functions while maintaining the modules original function.

A particular example in budding yeast’s cell-polarization module where duplication and sub-functionalization might have taken place is the diversification of the different GAPs of Cdc42 in budding yeast: Rga1, Rga2, Bem2, and Bem3: Bem3, Rga1, and Rga2 play individual roles in specific cellular functions, like the pheromone response pathway (Stevenson et al., 1995; Mukherjee et al., 2013), axial budding (Tong et al., 2007), and the timing of polarization (Knaus et al., 2007). This diversity of GAPs is promoted by cell-polarization mechanisms that are insensitive to GAP copy number, such as the Bem1-mediated WT mechanism. As we will argue below, this notion provides a concrete hypothesis about the role of scaffold proteins, like Bem1, for the evolution of functional modules that operate by the interplay of many interacting components.

### How evolution might leverage scaffold proteins

In the context of cellular signaling processes, it was suggested previously that evolution might leverage scaffold proteins to evolve new functions for ancestral proteins by regulating selectivity in pathways, shaping output behaviors, and achieving new responses from preexisting signaling components (Good et al., 2011). Our study of the Cdc42 polarization machinery shows how scaffold proteins may also play an important role in the evolution of intracellular self-organization. The scaffold protein Bem1 — by connecting Cdc42-GTP to Cdc42’s GEF — generates a functional submodule that contributes to selforganized Cdc42 polarization. Based on this, we propose a hypothetical evolutionary history for Bem1, illustrated in Figure 4: The latent rescue mechanism is generic and rudimentary as it requires only weak self-recruitment of Cdc42. The second requirement — enzyme saturation of Cdc42 hydrolysis — is a generic consequence of the enzymatic interaction between Cdc42 and its GAPs. That the same pattern-forming mechanism underlies MinD polarization in *E. coli* — based on the proteins MinD and MinE that are evolutionarily unrelated to the Cdc42 machinery — further underlines its generality. We therefore hypothesize that the latent rescue mechanism is a rudimentary, ancestral mechanism of Cdc42 polarization in fungi. On the basis of this ancestral mechanism, Bem1 could then have evolved in a step-wise fashion. Given that Bem1 is highly conserved in fungi (Diepeveen et al., 2018), and that fission yeast polarization is based on the same mutual recruitment mechanism (Lamas et al., 2019; Martin and Arkowitz, 2014), this hypothetical evolutionary pathway would likely lie far in the past.

How might step-wise evolution of Bem1 have occurred? A hypothetical Bem1 precursor binding to Cdc24 but not to Cdc42-GTP might have facilitated a globally enhanced catalytic activity of Cdc24 by relieving its auto-inhibition (Rapali et al., 2017; Shimada et al., 2004). Our theory shows that such an increase of GEF activity enlarges the range of GAP/Cdc42-concentration ratios for which the latent rescue mechanism is operational. This would have entailed an evolutionary advantage by increasing the robustness of the (hypothetical) ancestral mechanism against copy number variations. In a subsequent step the Bem1-precursor might then have gained the Cdc42-binding domain (SH3 domain) by domain fusion (Farr et al., 2017), thus forming the full scaffold protein that connects Cdc24 to Cdc42-GTP that mediates the WT polarization mechanism (mutual recruitment of Cdc24 and Cdc42). Along this hypothetical evolutionary trajectory, the constraints on the GAP/Cdc42 copy number ratio and the molecular properties of the GAPs (kinetic rates, membrane affinities) would be relaxed, thereby allowing the duplication and sub-functionalization of the GAPs (Conant and Wolfe, 2008).

There are several possible routes to test our hypotheses. One possibility is the construction of phylogenetic trees for the different proteins (domains) that could inform on the order they appeared during evolution of the polarity network (Hooff et al., 2019). Another possibility is to search for species in the current tree of life which contain intermediate steps of the evolutionary trajectory, for instance species with a more ancient version of Bem1 lacking the SH3 domain, and identify the protein self-organization principles underlying polarization in these species. This is becoming a more and more realistic option, given the very large (and still expanding) number of fungal species that has been sequenced (Diepeveen et al., 2018) and the growing interest of cell and molecular biologists to work with non-model systems (Russell et al., 2017).

Our mechanistic understanding of the polarization machinery provides a genotype–phenotype mapping where the molecular details have been coarse grained. In future work, one could integrate this map into a cell cycle model to address questions about epistasis, and eventually predict evolutionary trajectories in a population dynamics model.

On a broader perspective, we have shown how understanding the mechanistic principles underlying self-organization can provide insight into the evolution of cellular functions, a central theme in evolutionary cell biology. Specifically, we have presented a concrete example that shows how a self-organizing system can mechanistically evolve from more rudimentary, generic mechanisms that are parameter sensitive, to a specific, robust and tightly controlled mechanism by only incremental changes (Johnson and Lam, 2010).

## Materials and Methods

### Modeling and Theory

For theory materials and methods see the attached “Supplementary Information” file.

### Experiments

#### Media

All used media has the same base with 0.69% w/v Yeast nitrogen base (Sigma) + 0.32% Amino acid mix (4x CSM) (Formedium) + 2% Raffinose (Sigma), shortly, CSM+2% Raff. We used different galactose concentrations, denoted as *x*-Gal, where *x* denotes the Galactose percentage in media (*x* grams per 100 ml).

The plasmid pWKD010 contains P_gal_-Cdc42, URA3, Pre/Post-Cdc42 homology regions, with Ampicillin as a selectable marker on the pRL368 backbone (Wedlich-Soldner et al., 2004). The pWKD011 was based on pWKD010 but in this case a superfolder GFP (sfGFP) (Pédelacq et al., 2006) is integrated within in the CdC42 protein based on previous work in *S. cerevisiae*, where a mCherry was integrated within Cdc42 (Woods et al., 2015). We eliminated the fitness effects from mcherry-Cdc42^SW^ by using a superfolder GFP protein, as suggested by work in *S. pombe* (Bendezú et al., 2015). We comfirmed that the presence of the sfGFP insertion did not affect the growthrate of our cells in an detectable way in our growth rate assay compared to cells with Cdc42 under the Gal promotor without sfGFP.

#### Growth rate assays

We used a plate reader (Infinite M-200 pro, Tecan) for growth rate assays, with 96 well plates from Thermo Scientific, Nunc edge 2 96F CL, Nontreated SI lid, CAT.NO.: 267427. Rows A and H and the columns 1 and 12 were not used for measurements. We inoculated a 96-well plate with 100 μl of medium and 5 μl of cells (from glycerol stocks) in each well, and grew the cells in 96-well plate for 48 hours at 30 °C in a warm room. Afterwards the cells were diluted 200x into a new 96 well plate, which were then placed in the plate reader and the OD600 was measured for 48 hours using a combination of linear and orbital shaking. We used a home-written data analysis program in Matlab to determine the log-phase doubling time for every well. The doubling time was approximated by fitting the slope of the linear regime of the log plot of the raw data. We performed at least two different experiments per condition, and per experiments we performed at least 4 technical replicates per strain/plate.

The error in the growth rate plot is the 68 % credible interval of the posterior distribution of these rates. The posteriors of non-WT backgrounds followed from normalization to WT rates by Monte Carlo simulations of the quotient of the original, non-normalized growth rate posteriors in a genetic background and the WT posterior in that medium. The non-normalized posteriors were calculated using the Metropolis-Hastings algorithm (Hastings, 1970), from a rectangular prior and Student-t likelihood functions of doubling time fit estimates of all replicates in that medium. The standard errors of individual estimates come from the standard error of the slope parameter resulting from weighted least squares (WLS) on a moving window per OD curve, using an instrument error proxy for the WLS weights. The standard errors of individual estimates are corrected for overdispersion by the average modified Birge ratio (Bodnar and Elster, 2014) across media for WT.

#### Microscopy assays

All microscopy images were taken with an Olympus IX81 inverted microscope equipped with Andor revolution and Yokogawa CSU X1 modules. We used a 100x oil objective. The acquisition software installed is Andor iQ3. The CG imaging plates were from Zell-Kontakt. They are black multiwell plates compliant to the SBS (Society for Biomolecular Screening) standard-format with cover glass bottoms made from borosilicate glass.

Cells were grown in an overnight culture in CSM +2% Raffinose +2% Galactose media, without reaching saturation. On the next day, three washing steps with CSM+2% Raffinose were performed and subsequently the cells were re-suspended in the desired media of 0%, 0.06% and 0.1% Galactose. To obtain cell populations at all galactose concentrations, we first incubated all strains in 2% galactose concentration, where Cdc42 is highly overexpressed, such that also *bem1*Δ cells are able to efficiently polarize. After 15 hours of incubation in 2% galactose concentration, we exchanged the medium to the desired galactose concentration. After 24 hours, we observed the cells with light microscopy. After 24 hours leftover Cdc42 from the initial 2% galactose concentration incubation is (very low due to degradation and dilution (Cdc42 half-life is about 8 hours (Christiano et al., 2014)). From these images, we determined the average cell radius of the cells in the population.

Note that all of them contain the same base media: CSM+2% Raffinose. Afterwards the cells were incubated for 8 hours at 30°C, followed by an imaging session, and subsequently incubated for another 16 hours after which another imaging sessions was performed. We performed three independent experiments for each galactose concentration.

#### Microscopy data analysis

We performed bright field microscopy assays to monitor the cell size across different levels of Cdc42 in different genetic backgrounds. With ImageJ we manually determined the perimeter of the individual cells by fitting the live cells to a circle with the Measure tool. We performed three independent experiments per condition and per strain. In addition, we visually checked how many of the cells were alive and how many were dead based on their morphology. The error bar on the fraction of dead cells as well as of the average cell radius, is calculated as the standard error over the total number of analysed cells.

**Table 1.** Strains used in this work

| Name | Genotype | Source |
| --- | --- | --- |
| yLL3a | *MATa can1-100, leu2-3, 112, his3-11,15, ura3Δ, BUD4-S288C* | (Laan et al., 2015) |
| yWKD065a | *MATa ,*<br>*-P<sub>gal</sub>-sfGFP-Cdc42<sup>SW</sup> (pWKD011 integrated),*<br>*leu2-3, 112, his3-11,15, ura3Δ, BUD4-S288C* | This work |
| yWKD069a,b,c | *MATa, bem1:: KanMX6, URA-P<sub>gal</sub>-sfGFP-Cdc42<sup>SW</sup> (pWKD011 integrated)*<br>*can1::P<sub>mfa</sub>-HIS3 1, leu2-3, 112, his3-11,15, ura3Δ, BUD4-S288C* | This work |
| yYWKD070a | *MATa, bem1:: KanMX6, bem3::NATMX4, URA-P<sub>gal</sub>-sfGFP-Cdc42<sup>SW</sup> (pWKD011 integrated), can1:: P<sub>mfa</sub>-HIS3, leu2-3, 112, his3-11,15, ura3Δ, BUD4-S288C* | This work |
| yYWKD071a | *MATa,*<br>*URA3 P<sub>gal</sub>- CDC42 (pWKD010 integrated),*<br>*can1:: P<sub>mfa</sub>-HIS3, leu2-3, 112, his3-11,15, ura3Δ, BUD4-S288C* | This work |
| yYWKD073a | *MATa, bem1:: KanMX6, bem3::NATMX4, URA3- P<sub>gal</sub>-CDC42 (pWKD010 integrated), can1:: P<sub>mfa</sub>-HIS3, leu2-3, 112, his3-11,15, ura3Δ, BUD4-S288C* | This work |

## Supporting information

Simulation Videos

## Supplementary Information

## 1 Model description

The primary goal of the mathematical model we propose is to explain the rescue of *bem1*Δ. cells by the loss of *BEM3*. To that end, we make minimal, but essential extensions to a previously established model (Klünder et al., 2013) that accounts for the core Cdc42-polarization mechanism relying on the Bem1-mediated pathway (Irazoqui et al., 2003; Kozubowski et al., 2008; Bendezú et al., 2015; Witte et al., 2017; Chiou et al., 2017). Importantly, the extended model we propose here enables us to explain several previous experimental findings that had remained puzzling so far. We will summarize and discuss these findings that serve as additional support for our model in the supplementary discussion (Sec. 6.1).

In what follows, we describe the biophysical and biochemical processes (diffusion, vesicle-based transport and protein interactions) accounted for by our model. The mathematical formulation and analysis of the model in the framework of bulk-surface coupled reaction–diffusion systems is presented in the subsequent sections.

### 1.1 Transport processes and protein interactions

The mathematical model we propose is based on the quantitative model introduced in (Klünder et al., 2013), which we extended in several important ways. The cell is modeled as a spherical domain with a diffusive bulk (cytosol) on the inside and the membrane on the surface where proteins interact and diffuse laterally. Bulk and surface dynamics are coupled due to membrane attachment and detachment of proteins. Mathematically, the model is formulated as a reaction–diffusion system with bulk-surface coupling (see Sec. 2.1). As we will argue in the following, both transport pathways — cytosolic cycling and vesicle-based transport (Figs. 1B and 1C in the main text) — can be incorporated in this modeling framework.

In previous works, vesicle-trafficking along actin cables has been modeled to various degrees of detail and based on different assumptions (Wedlich-Soldner, 2003; Slaughter et al., 2009; Hawkins et al., 2009; Layton et al., 2011; Savage et al., 2012; Freisinger et al., 2013; Muller et al., 2016). However, a mechanistically detailed modeling of vesicle trafficking is not feasible at the moment because the highly complex vesicle recycling pathway — involving endocytosis, transport along actin cables, processing in intracellular membrane compartments like endosomes and the Golgi apparatus, and finally exocytosis — is not fully characterized experimentally. As we will see, however, a detailed description is not required for the purpose of the analysis here. Instead, we model vesicle recycling of Cdc42 as effective membrane-recruitment of Cdc42-GDP by Cdc42-GTP. This effective description incorporates the two essential features of vesicle recycling that are relevant for the polarization machinery: (*i*) vesicle transport is directed towards membrane-bound Cdc42-GTP and (*ii*) Cdc42 delivered to the membrane by vesicles upon exocytosis is (mostly) GDP-bound (Savage et al., 2012). Details are discussed in Sec. 1.2.

In addition to vesicle recycling, several downstream effectors of Cdc42-GTP — Cla4, Gic1/Gic2, and flippases (Brown et al., 1997; Chen et al., 1997, 2012; Das et al., 2012; Tiedje et al., 2008) — have been suggested to facilitate membrane-recruitment of Cdc42-GDP (see Sec. 1.3 for details). We incorporate these putative Cdc42-GDP-recruitment pathways together with vesicle-based Cdc42-transport by a single, effective recruitment process that is directed by membrane-bound Cdc42-GTP (illustrated in Fig. 1D (4) in the main text).

Figure 1D shows the biochemical interaction network underlying our model. At its core is the GTPase cycle of Cdc42 ((1) in Fig. 1D). Cdc42 cycles between an active, GTP-bound, and an inactive, GDP-bound state on the membrane. In its GDP-bound form, Cdc42 can bind to the guanine-nucleotide-dissociation inhibitor (GDI) Rdi1, which sequesters Cdc42’s membrane binding anchor and thus enables it to diffusive freely in the cytosol. The cycling of Cdc42 between its GTP- and GDP-bound states is regulated by the guanine nucleotide-exchange factor (GEF) Cdc24, and GTPase-activating proteins (GAPs) that catalyze the hydrolysis from GDP to GTP. In wild-type cells, Cdc42-GTP recruits the scaffold protein Bem1 to the membrane which in turn recruits the GEF Cdc24 ((2) in Fig. 1D) to form a Bem1–GEF complex. These membrane-bound Bem1–GEF complexes recruit Cdc42-GDP from the cytosol to the membrane and activate it there ((3) in Fig. 1D), thus closing the feedback loop (mutual recruitment) that underlies WT polarity (Bose et al., 2001; Irazoqui et al., 2003; Goryachev and Pokhilko, 2008); see Chiou et al. (2017) and Halatek et al. (2018) for recent reviews through the experimental and theoretical lens, respectively.

Extending previous models (Goryachev and Pokhilko, 2008; Klünder et al., 2013), we explicitly incorporate transient formation of a Cdc42-GAP complex as an intermediate step in the enzymatic interaction between GAPs and Cdc42 (Zhang et al., 1997). This explicit modeling of the enzymatic reaction dynamics is important if the dissociation of the substrate-enzyme (here, Cdc42-GAP) complex is the rate limiting step. Indeed, it has been shown that this is the case for GAP-catalyzed hydrolysis of Cdc42 in budding yeast (Zhang et al., 1997).

In budding yeast, four GAPs for Cdc42 are known: Bem2, Bem3, Rga1, and Rga2 (Bi and Park, 2012). It has been found that the different GAPs have specific roles in several cellular functions coordinated by Cdc42 polarization, such as pheromone response (Stevenson et al., 1995; Mukherjee et al., 2013), axial budding (Tong et al., 2007), and the timing of bud emergence (Knaus et al., 2007). For the purpose of our model, we disregard their differences and conflate them into a single effective GAP-species. We account for cycling of GAP proteins between two states: free and Cdc42-bound, both on the membrane. Importantly, in our model, the GAP protein copy number is an explicit parameter and the Bem3 deletion is accounted for by decreasing the copy number of Bem3. Incorporating more states of the GAPs (e.g. a cytosolic state, phosphorylation, etc.) and shuttling of GAPs to internal membranes (Mukherjee et al., 2013) is not required to explain the rescue pathway for *bem1*Δ. cells and goes beyond the scope of the present work.

Because Cdc24 has its own membrane-binding (PH) domain (Rossman et al., 2005), we also incorporate Cdc24 membrane binding independently of Bem1. Compared to Cdc24 in a complex with Bem1, free membrane-bound Cdc24 has an approximately 50% lower GEF activity towards membrane-bound Cdc42 (Rapali et al., 2017; Shimada et al., 2004). We assume that, in the absence of Bem1, Cdc24 will have linear attachment–detachment kinetics without feedback such that the membrane-bound fraction of Cdc24 will distribute uniformly.

In summary, the extensions of the model introduced in (Klünder et al., 2013) are:

1. Explicit modeling of Cdc42’s hydrolysis by GAPs as a catalytic reaction with an intermediate Cdc42-GAP complex (Zhang et al., 1997).
2. Effective membrane-recruitment by membrane-bound Cdc42-GTP, accounting for vesicle-based transport of Cdc42 towards zones of high Cdc42-GTP concentration as well as further putative recruitment pathways mediated by downstream effectors of Cdc42-GTP (Tiedje et al., 2008; Das et al., 2012; Daniels et al., 2018; Grinhagens et al., 2020).
3. Membrane binding of the GEF Cdc24 independently of Bem1 via Cdc24’s PH domain (Rossman et al., 2005).

The model analysis in terms of functional subunits performed in the main text shows that all three extensions are required to describe the rescue of *bem1*Δ. mutants in the model. Let us conclude the model description with several additional remarks:

- A second (Bem1-independent) positive feedback loop for the GEF (Cdc24) has been hypothesized in previous literature (Witte et al., 2017; Goryachev and Leda, 2017). We do not incorporate such a feedback loop in our model. Moreover, we show in Sec. 6.1 that our model can provide an explanation for the experimental findings of (Witte et al., 2017) without the hypothesized second GEF feedback loop. If they are present, such feedback loops would impart additional redundancy on the Cdc42 polarization machinery and thus add to the robustness of this machinery against genetic perturbations. Identifying these feedback loops in experiments and accounting for them in mathematical models would be an interesting future extension of our work.
- We use the protein copy numbers for Cdc42, Bem1, Cdc24 and the GAPs reported in (Kulak et al., 2014) to ensure that the values have been obtained by the same method (termed “in-StageTip”) and thus are consistent relative to one another. This consistency is important because the relative ratios of the protein copy numbers are what matters for operation of the various Cdc42 polarization mechanisms. In contrast, the earlier study (Klünder et al., 2013), used values from a range of several distinct, older sources. Note that there are some significant differences: The copy numbers of Cdc42 and Bem1 reported in (Kulak et al., 2014) are larger by factors 3 and 6 compared to the values used in (Klünder et al., 2013).
- As the earlier model from (Klünder et al., 2013), our model includes direct recruitment and activation of Cdc42 from the cytosol to the membrane. Other models (e.g. (Kuo et al., 2014)) don’t assume this and instead only incorporate activation of Cdc42 on the membrane by Bem1-GEF complexes. Both types of models are qualitatively identical as both capture the key effect that Bem1-GEF generates a sink for Cdc42-GDP and thus leads to directed (cytosolic) diffusion of Cdc42-GDP towards the polar zone.
- For simplicity, we disregard slow intrinsic nucleotide exchange and intrinsic hydrolysis of Cdc42 (which both were reported on the order of 10^−3^ s^−1^, see Zheng et al. (1994); Zhang et al. (1997)). As these processes are slow and do not impart nonlinear feedback, including these processes does not change our results qualitatively.
- The model does not incorporate the interaction of the Cdc42 polarization machinery with upstream cues (landmark proteins, pheromone signals) that are important for timing of polarization, bud-site selection (see e.g. Miller et al. (2017); Kang et al. (2014, 2018); Miller et al. (2019)), and shmoo formation (Muller et al., 2016). For conceptual cell-polarity models with two components, it was shown in a recent publication (Brauns et al., 2018) that the ability to exhibit spontaneous polarization is a necessary requirement for the maintenance of stationary patterns that are induced by a spatial cue. We therefore expect that our results may also be relevant for the ability of the Cdc42 polarization machinery to exhibit cue-guided polarization.

### 1.2 Remarks on vesicle-based transport

Vesicles are transported towards exocytosis sites along polarized actin cables. In budding yeast, the formation of such actin cables is induced by the formin Bni1 which in turn is recruited by Cdc42-GTP. The collective effect of these processes lead to directed transport of vesicle-bound proteins (Cdc42, for instance) to membrane sites of high Cdc42-GTP concentration. Moreover, it was found that Cdc42-GTP activates the exocyst tethering complex and thus promotes vesicle exocytosis in the polar zone independent of actin cables (Wu et al., 2008). However, the role of vesicle-based Cdc42 transport for cell polarization remains under debate (Howell et al., 2009; Layton et al., 2011; Watson et al., 2014; Woods et al., 2016).

Vesicle recycling might also indirectly promote Cdc42 polarity. First, it was found that specific lipids (phosphatidylserine) are transported to the polar zone by vesicles and that these lipids in turn promote Cdc42 clustering, activation and membrane-binding (Fairn et al., 2011). This might result in Cdc42 transport to the polar zone via cytosolic transport and by trapping Cdc42 (see Sec. 1.3). Second, vesicle recycling has been shown to be important for septin-ring formation by diluting septins in the center of the polar zone (Okada et al., 2013). Because septins recruit the GAP Bem2, this effectively transports Bem2 away from the center of the polar zone. Thus, vesicle recycling might have a stabilizing effect on Cdc42 polarity by reducing Cdc42 deactivation within the polar zone (Martin, 2015).

In our reaction–diffusion model, we incorporate directed transport by vesicle recycling as an effective recruitment process of cytosolic Cdc42-GDP to the membrane. This may seem counterintuitive at first. On closer inspection, however, it turns out that both processes share the same key features that are relevant for the Cdc42-polarization machinery. In fact it ha been shown experimentally, that the cytosolic transport directed by membrane recruitment and vesicle-based transport directed by formins and the exocyst complex are functionally interchangeable:

- A Bem1-Snc2 fusion chimera, which is permanently membrane bound and transported on vesicles, rescues *bem1*Δ. rsr16. mutants (Howell et al., 2009; Chen et al., 2012). This shows that diffusive transport of Bem1 to the polar zone due to recruitment by Cdc42 can be replaced by transport on vesicles along directed actin cables (“vesicle recycling”).
- A Cdc24^PBΔ^-Snc2 fusion chimera which does not interact with Bem1 and is transported on vesicles completely bypasses the Bem1-mediated recruitment pathway for Cdc24 (Woods et al., 2015). In the same study, it is shown that a Cdc24^PB.6.^-Cla4 fusion chimera that is directly recruited to Cdc42-GTP bypasses Bem1-mediated recruitment as well. In their summary, the authors explicitly point out the interchangeability of transport pathways: *“Thus, the functional deficit of a Cdc24 that lacks the PB1 domain can be rescued by linkage to a polarized protein, whether that protein polarizes by diffusion capture (Cla4) or vesicle recycling (Snc2).*”

Underlying the functional interchangeability of cytosolic transport and vesicle-based transport in the Cdc42-polarization machinery is that both are directed by Cdc42-GTP, as illustrated in Fig. 1B,C in the main text. Hence, in a *coarse-grained* description, vesicle-based transport and cytosolic diffusion directed by membrane-recruitment are equivalent and can be modeled in the framework of bulk-surface coupled reaction– diffusion dynamics.

### 1.3 Putative recruitment of Cdc42 to the polar zone

There is some experimental evidence that several downstream effectors of Cdc42-GTP, such as Cla4, Gic1/2, and flippases, may mediate membrane-recruitment of Cdc42-GDP. These recruitment mechanisms are likely to drive Cdc42 transport towards the polar zone independently of vesicle-based transport and independently of Bem1: Cla4 has been found to catalyze the release of Cdc42-GDP from its GDI (Rdi1) (Tiedje et al., 2008). Gic1/2 have been found to stabilize Cdc42-GTP on the membrane, reducing detachment and lateral diffusion away from the polar zone (Bendezú et al., 2015; Daniels et al., 2018; Kang et al., 2018). Flippases downstream of Cdc42 flip specific phospholipids to the inner membrane leaflet. These phospholipids have been reported to decrease the detachment of Cdc42 (Das et al., 2012). Moreover, phosphatidylethanolamine, a phospholipid that promotes Cdc42-extraction from the membrane by Rdi1 is flipped to the outer leaflet of the membrane in the polar zone (Das et al., 2012). This effectively reduces Cdc42 detachment in the polar zone compared to the remaining membrane surface.

While the molecular details and functional relationships remain elusive, several experimental findings provide indirect evidence for these putative self-recruitment pathways of Cdc42. First, an overexpression of Gic1 but not of Gic2 was found to partially rescue *bem1*Δ cells (Brown et al., 1997). Conversely, it was recently found that over-expression of Gic2 is lethal for *bem1*Δ *bem3Δ* cells (Grinhagens et al., 2020). Second, *gic1Δ gic2Δ cla4Δ* triple-mutants were reported to grow very slowly (Chen et al., 1997). Moreover, this study showed that *gic1Δ gic2Δ*. mutants are highly temperature sensitive (strongly impaired growth at 35 °C) and that overexpression of Cla4 can suppress the growth defect of *gic1Δ gic2Δ* mutants at 35 °C. Finally, also overexpression of Bem1 or of Cdc42 in its WT form, but not in the GTP-locked state (G12V), were found to suppress the growth defect of *gic1*Δ. *gic2Δ* mutants at 37 °C (Brown et al., 1997).

A recent study established that the growth defects in *gic1Δ gic2Δ* mutants at high temperature are in fact due to impaired Cdc42-polarization (Daniels et al., 2018). Strikingly, in this study it was also found that *gic1Δ gic2Δ* cells at 37 °C quickly assume a rescue mutation that restores the wild-type behavior. This mutation reproducibly happens at a single locus, its exact identity remains unknown though (Daniels et al., 2018).

Together, these experimental findings suggest that Gic1/2 and Cla4 provide several, potentially redundant pathways of effective Cdc42 self-recruitment feedback loops that become relevant at high temperature to “support” or entirely replace the Bem1-mediated polarization mechanism. In our model, such feedback loops are accounted for by the generic self-recruitment term with rate *k_tD_* Studying the details of these feedback mechanisms remains an open task for experimental and theoretical studies.

## 2 Reaction–diffusion dynamics with bulk-surface coupling

### 2.1 General framework

Since budding yeast cells are (nearly) spherical, we study the proteins’ reaction– diffusion dynamics in a spherical geometry composed of a cytosol (bulk) of radius R with membrane on its surface (Fig. S1). Naturally, we choose spherical coordinates (*r*, φ, θ). For a general, compact notation, we denote concentrations of membrane-bound and cytosolic components by vectors **m** and **c**, respectively.

In the bulk, we consider purely diffusive dynamics,

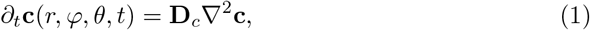

with the matrix of diffusion constants **D**_*c*_ = diag({D_*i*_}). Unless stated otherwise, the cytosolic diffusion constants are all set to the same value D_*c*_ such that **D** _*c*_ = D_*c*_. In spherical coordinates, the Laplacian ∇^2^ acting on some ψ function reads

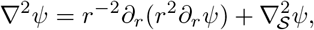

where the “angular” Laplacian on the sphere’s surface 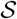 is given by

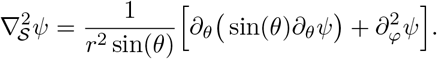

The bulk is coupled to the membrane by attachment-detachment reactions that lead to bulk flows, ***f***, normal to the surface

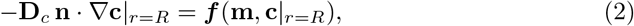

where **n** is the surface’s inward normal vector. In spherical coordinates, the radial gradient is given by the radial derivative **n**. ∇ = −∂_*r*_. The attachment-detachment flows ***f*** of our specific model will be specified further below in Eq. (4).

**Figure S1.**
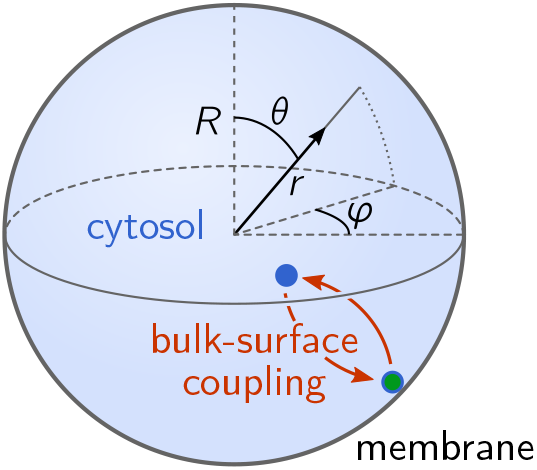
Spherical cell geometry with spherical coordinates (*r, φ, θ*) and an illus-tration of the bulk-surface coupling due to attachment–detachment dynamics at the membrane.

The dynamics of membrane-bound components are given by

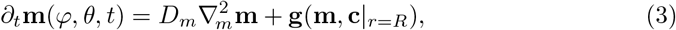

where the nonlinear function **g** encodes the nonlinear reactions on the membrane. Note that the diffusion operator on the membrane 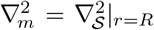 coincides with the bulk Laplacian ∇^2^ restricted to the membrane at *r* = *R*. This is because the sphere fulfills the rotational symmetries of the diffusion operator.

### 2.2 Variables, reaction terms, and conserved protein numbers

As shorthands for the protein concentrations we use the same shorthand notation as in Fig. 1 in the paper: D – Cdc42-GDP; T – Cdc42-GTP; G – GAPs; B – Bem1; F – GEF. (Note that we refer to Cdc24 as GEF to prevent confusion with Cdc42.) We denote concentrations of membrane-bound species with the symbol *m* with lowercase subindices, and cytosolic concentrations using the symbol *c* with uppercase subindices (see Table S1). Using the vector notation introduced above, we have **c** = (*c*_D_, *c*_B_, *c*_F_), **m** = (*m*_d_, *m*_t_, *m*_tg_, *m*_g_, *m*_b_, *m*_bf_, *m*_f_).

The protein interactions described above in Sec. 1 and illustrated in Figure 1 in the main text are modeled by mass-action law kinetics, with the reaction rates described in Table S2:

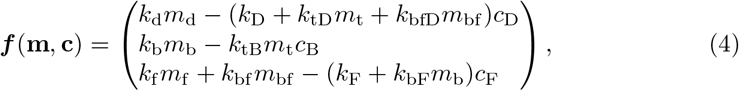

and

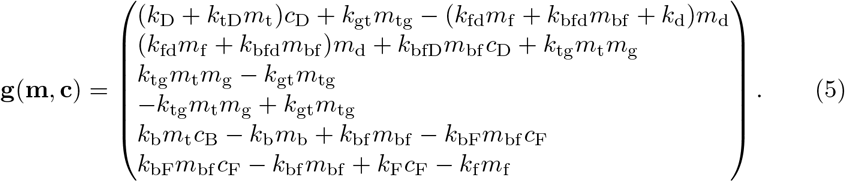

**Table S1.** Variables of the reaction–diffusion model describing the protein concentrations of Cdc42, GAPs, Bem1 and GEF (Cdc24) in various conformational states — cytosolic, membrane bound and in form of multi-protein complexes.

| Domain [Unit] | Symbol | Description |
| --- | --- | --- |
| Cytosol [$\mu\text{m}^{-3}$] | $c_D$ | Cdc42-GDP (potentially GDI-bound) |
| | $c_B$ | Bem1 |
| | $c_F$ | GEF |
| Membrane [$\mu\text{m}^{-2}$] | $m_d$ | Free Cdc42-GDP |
| | $m_t$ | Cdc42-GTP |
| | $m_g$ | GAP |
| | $m_{tg}$ | Heterodimeric Cdc42-GAP complexes |
| | $m_b$ | Bem1 |
| | $m_{bf}$ | Heterodimeric Bem1-GEF complexes |
| | $m_f$ | GEF |

These dynamics conserve the total numbers of Cdc42, GAPs, Bem1, and GEF molecules,

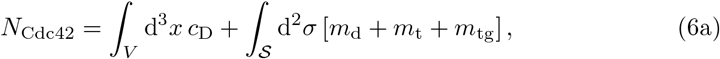

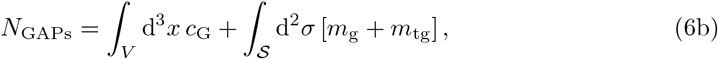

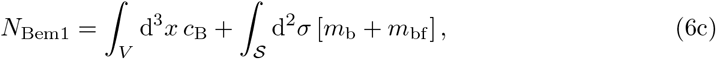

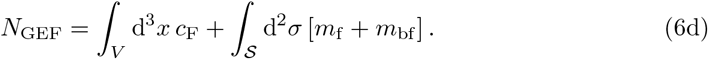

Hence, these protein copy numbers are control parameters of the model.

**Table S2.**
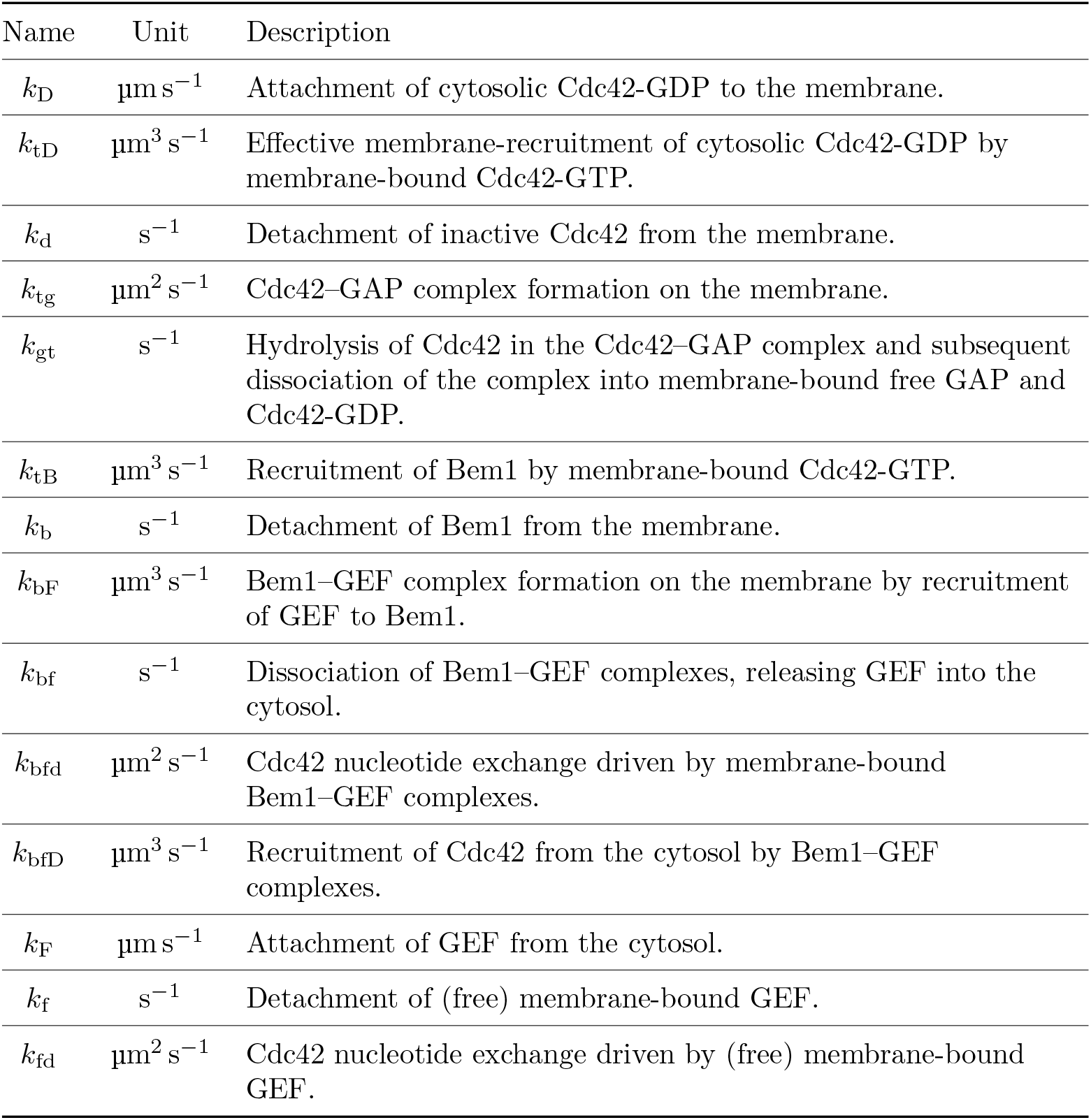
Reaction rates with descriptions. For the parameter values used to exemplify the results and a detailed discussion how these values were chosen see Sec. 5.

## 3 Linear stability analysis in spherical geometry

### 3.1 Laterally homogeneous steady states

Our goal is to determine the linear stability of steady states that are homogeneous on and at the membranes: 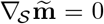 and 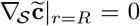. Because the bulk dynamics are purely diffusive, the radial bulk profiles corresponding to these steady states are also homogeneous, that is, 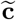 is constant in the entire bulk. The homogeneous steady states are thus determined by the set of equations

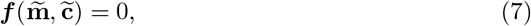

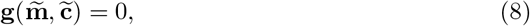

together with the total density constraints

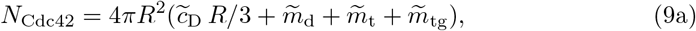

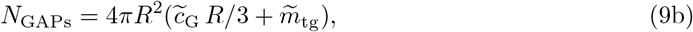

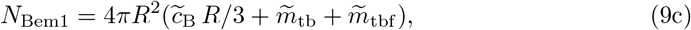

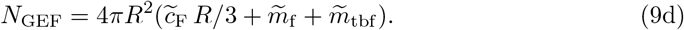

We solve this set of algebraic equations numerically (implementation in Mathematica using the built-in function NSolve[]).

### 3.2 Linearized dynamics

Linear stability of a steady state is studied by calculating the growth rate of small perturbations (δ**m**, δ**c**) around the steady state. For sufficiently small perturbations, the dynamics can be linearized

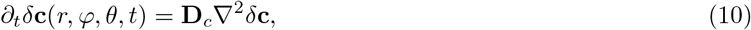

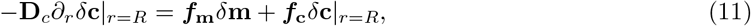

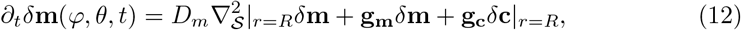

where the matrices ***f_c, m_*** and **g_c_** _**m**_ are the linearized attachment-detachment kinetics and membrane reactions,

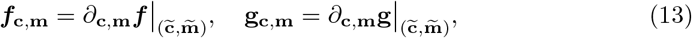

evaluated at the steady state 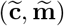. In the case of a homogeneous steady state, these matrix coefficients are constant in space. This allows us to find the spatial eigenmodes of the linearized dynamics analytically and reduce the set of linear PDEs (10)–(12) to an algebraic problem that then can be solved numerically.

### 3.3 Spatial eigenmodes and finding their growth rates

The key idea to solve the linearized dynamics Eqs. (10)–(12) is to consider elementary perturbations of the form (Levine and Rappel, 2005; Klünder et al., 2013)

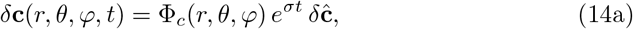

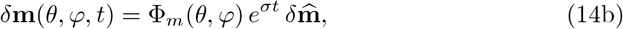

where Φ_*c,r*_ are the spatial eigenmodes that encode the spatial form of the perturbation and CJ is its temporal growth rate. The goal is to find triplets (σ, Φ_*c*_, Φ_*m*_) that fulfill Eqs. (10)–(12). In principle, a general solution of the linearized dynamics can be constructed from a superposition of elementary perturbations. However, we are mostly interested in the stability of the steady state which is determined by the growth rate with the largest real part. If it is positive, the steady state is linearly unstable because the corresponding elementary perturbation will grow exponentially in time.

To solve the linear PDEs (10)–(12), the ansatz Eq. (14) requires that the spatial eigenmodes Φ_*m*_(*θ*, *φ*), Φ_*c*_(*θ*, *φ*, *r*) simultaneously diagonalizes all spatial derivative operators encoding (i) diffusion in the bulk, (ii) diffusion on the membrane, and (iii) the bulk-boundary coupling. Because the spherical geometry obeys the rotational symmetries of the diffusion operators in the bulk and on the surface, we can find such spatial eigenmodes by a separation of variables

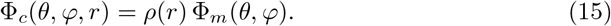

Inserting this ansatz into the diffusive bulk dynamics yields^1^

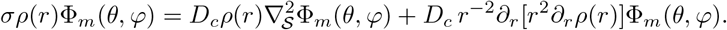

Because there are no mixed derivatives, we can rewrite the previous equation as

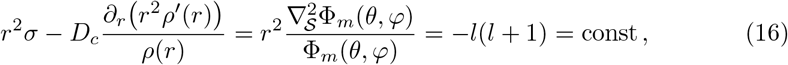

where we introduced the separation constant −*l*(*l*+1) such that the angular eigenmodes are the spherical harmonics 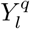 that solve

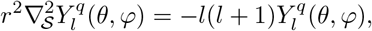

where 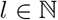 and 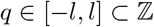 are the ‘degree’ and ‘order’ of the spherical harmonic (Bronshtein, 2007). Essentially, the degree determines the number of peaks in the function 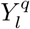, whereas the order q determines where these peaks are located on the sphere. Due to the system’s rotational symmetry, only the degree l enters in the linear stability problem. The mode *l* = 0 is uniform, modes with *l* = 1 correspond to a polar patterns, modes with *l* = 2 correspond to bi-polar patterns and so on.

For the radial bulk profile *ρ*(*r*), Eq. (16) mandates

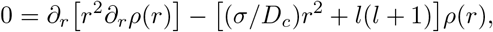

which, using the rescaled radius 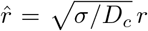, is identical to the modified spherical Bessel equation, which are solved by the modified spherical Bessel functions of the first kind *i_*l*_*(*r*) (Bronshtein, 2007). (The spherical Bessel functions of the second kind diverge at *r* = 0 and are therefore unphysical in the setting considered here.) With the convenient normalization *ρ*(*R*) = 1, the radial bulk profiles are given by

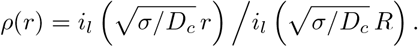

Substituting this bulk solution with the ansatz Eq. (14) into the linearized bulk-surface coupling Eq. (11) and membrane dynamics Eq. (12) yields

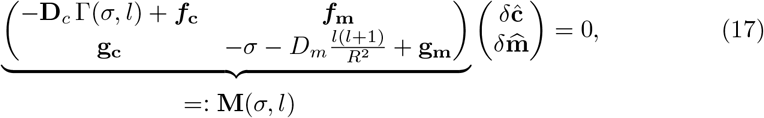

with the bulk-surface coupling coefficient

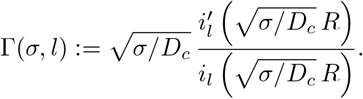

Using the relationship 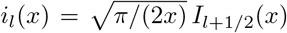, where *I_*k*_*(*x*) denotes the modified Bessel functions of the first kind, one can further evaluate Γ to

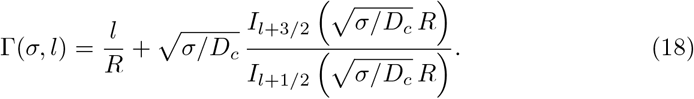

In the general case of different cytosolic diffusion constants, *D*_*c,i*_ (index i denoting cytosolic components), the bulk-surface coupling coefficient is a diagonal matrix **Γ** = diag({Γ_*i*_}) where D_*c*_ is replaced the respective component’s bulk diffusivity D_*c,i*_ in each Γ_*i*_.

The system of linear equations Eq. (17) has non-trivial solutions 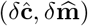 only when the determinant of the matrix **M** (σ, *l*) vanishes. Hence, the growth rates σ_*l*_ of spatial perturbations with the degree *l* are determined by the complex solutions of the solvability condition (characteristic equation)

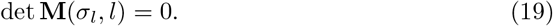

The stability of a spatial perturbation is determined by the respective growth rate σ_*l*_ with the largest real part. We solve this problem numerically using the iterative Newton method (implementation in Mathematica, see supplementary file `LSA-setup.nb`).

**Figure S2.**
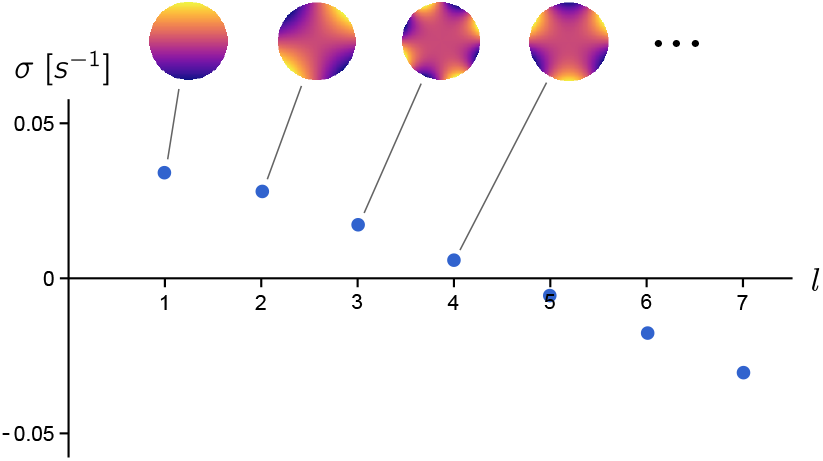
Example for a dispersion relation in spherical geometry, showing the growth rate CJ as a function of the spherical harmonic order *l*. The density plots above illustrate the spherical harmonics with *l* = 1…4.

An example of the resulting relation between the spherical harmonic order *l* and the dominant growth rate σ_*l*_ is shown in Fig. S2. We have used this linear stability analysis to identify the parameter regimes that exhibit spontaneous Cdc42 polarization, i.e. where the homogeneous steady state is laterally unstable (see Sec. 5 below and the stability diagrams in Figs. 2, 4 and 5 in the main text).

## 4 Numerical simulations

To validate our findings from linear stability analysis and determine the eventual steady state patterns formed, we performed numerical simulations in COMSOL Multiphysics. This software uses Finite Element Methods (FEM) and has out of the box support for systems with bulk-surface coupling. Simulation results for the various mutations and conditions that were studied experimentally are shown in Videos 1–6. The simulations are initialized with Cdc42, GEF and Bem1 distributed uniformly in the cytosol, with a small random perturbation added (5% random noise). The GAP density is initialized uniformly on the membrane. The parameters used for the numerical simulations are given in Tables S3 and S5, where the former table contains all parameters that have been directly determined in experiments and the latter table states the kinetic rates that were determined by parameter sampling as described in Sec. 5.

In addition, we performed a numerical simulation that emulates the optogenetic recruitment of Cdc24 to a spot on the membrane in *bem1*Δ. cells (Witte et al., 2017). The optogenetic recruitment is implemented via a spatially dependent increase of Cdc24-attachment rate to 1.0 μm s^−1^ in the shape of a Gaussian pulse with a radius of 1 μm starting at 500 s and ending at 1000 s in the simulation. In agreement with the experiments, we find that this transient, localized GEF recruitment stimulates Cdc42-polarization and that this polarization is maintained after the localized GEF recruitment is switched off (see Video 6).

## 5 Inference of reaction rate parameters

In the main text, we reasoned why the the rescue mechanism generically is operational only when the GAP to Cdc42 copy number ratio is below a threshold. The quantitative value of the threshold depends on the parameters in the model, many of which are not directly constrained by experiments. However, because we know from previous experiments that polarization is impaired in *bem1*Δ. but operational in *bem1*Δ. bem36. (Laan et al., 2015), we can put bounds on the critical GAP-Cdc42 ratio from above and below: Based on the protein copy numbers reported in Kulak et al. (2014), cf. Table S3, the critical ratio lies above (*N*_GAPs_ −*N*_Bem3_)/*N*_Cdc42_ ≈ 0.18 and below *N*_GAPs_/*N*_Cdc42_ 0.25. Based on this estimate directly from previous experiments, we can make predictions on the effect of changing the Cdc42 copy number (via a GAL promoter) on different mutant strains, without the need to specify parameter values for the model.

We still need to show that the experimentally found critical GAP-Cdc42 ratio is actually exhibited by the mathematical model for physiologically realistic parameter values. To that end, we fix those parameters that are directly constrained by experimental measurements, and perform massive random sampling of the remaining parameters, filtering for those parameter sets that are consistent with our experimental findings on Cdc42 copy number dependence of cell polarity of WT, *bem1*Δ. and *bem1*Δ.*bem3*Δ. cells, as well as the previous experimental observation that a permanently membrane-bound Cdc42 mutant (Cdc42-rit^C^) is able to polarize (Bendezú et al., 2015; Kang et al., 2018). The technical details of this parameter *sampling procedure* are provided in Sampling procedure the subsequent paragraphs below. The Mathamtica code implementing this procedure is available in the supplementary file `parameter-filtering.nb`.

Importantly, we find that almost all parameter can be varied over the entire sampled range (four orders of magnitude) indicating that the model is sloppy (Gutenkunst et al., 2007; Daniels et al., 2008). For such models, the collective behavior can usually not be used to infer (tightly constrain) the underlying parameters (Transtrum et al., 2010). From the parameter sets identified by the large scale sampling, we picked one representative example to generate the stability diagrams shown in the main text (Figs. 2 and 5). In addition, we filtered under the extra condition of permanently membrane-bound, immobilized Cdc42 to demonstrate that a polarization mechanism based on Bem1–GEF redistribution and local cycling of Cdc42 between active and inactive states (Fig. 4H) can coexist in overlapping parameter regimes with the other two polarization mechanisms (WT and rescue). A representative example out of the parameter sets obtained with this additional condition was used to generate the stability diagrams shown in Fig. 5C-E.

### Sampling procedure

Each of the 13 reaction rates (see Tab. S2) is sampled over four orders of magnitude from 10^−3^ to 10^1^ uniformly on a logarithmic scale. To allow for exact reproduction of the random sampling, the random number generator in Mathematica is seeded with a specified number. For each parameter set, the homogeneous steady state and its dispersion relation (linear stability) are computed under various conditions that emulate the mutants that were studied experimentally (see Tab. S4).

**Table S3.** Parameters based on literature values (directly determined by experiments). Note that the protein copy numbers are stated for wild-type (W)T cells. To account for genetic perturbations (mutations), the values are adapted as specified in Table S4.

| Parameter | Value | Description / Comments |
| --- | --- | --- |
| $R$ | $3.5 \mu\text{m}$ | Cell radius. For the parameter sampling, we checked that larger cell size, $R = 7 \mu\text{m}$, alone does *not* rescue polarization of *bem1*Δ cells. |
| $N_{\text{Cdc42}}$ | 8690 | Cdc42 copy number (WT) (Kulak et al., 2014). |
| $N_{\text{Bem1}}$ | 1037 | Bem1 copy number (WT) (Kulak et al., 2014). |
| $N_{\text{GEF}}$ | 934 | Cdc24 copy number (WT) (Kulak et al., 2014). |
| $N_{\text{GAPs}}$ | 2170 | Aggregate copy number of all four Cdc42-GAPs. The individual numbers are 117, 332, 1122, and 599 for Rga1, Rga2, Bem2, and Bem3, respectively (Kulak et al., 2014). |
| $D_c$ | $11 \mu\text{m}^2 \text{s}^{-1}$ | Cytosolic diffusion constants (Slaughter et al., 2007). |
| $D_{m_d}$ | $0.2 \mu\text{m}^2 \text{s}^{-1}$ | Diffusion of membrane-bound Cdc42-GDP (Bendezú et al., 2015). |
| $D_{m_t}, D_{m_{tg}},$<br>$D_{m_g}, D_{m_b},$<br>$D_{m_{bf}}, D_{m_f}$ | $0.01 \mu\text{m}^2 \text{s}^{-1}$ | Diffusion of the remaining membrane-bound proteins. There is no consensus on the membrane diffusion rates in the literature. We use a value that lies in-between the two values $0.03 \mu\text{m}^2 \text{s}^{-1}$, and $0.0025 \mu\text{m}^2 \text{s}^{-1}$ used in previous models (Klunder et al., 2013; Kuo et al., 2014; Goryachev and Pokhilko, 2008). |
| $k_{\text{bfd}}$ | $2 \cdot k_{\text{fd}}$ | The GEF activity of Bem1-Cdc24 complexes increased approx. two-fold compared to the GEF activity of Cdc24 alone (Shimada et al., 2004). |

**Table S4.** Conditions for the filtering of randomly sampled parameters based on experimental findings from this study (1–6, highlighted in bold) and the previous works (Bendezú et al., 2015; Kang et al., 2018) for Cdc42-rit^C^ (8) and Cdc42-psy1™ in *S. pombe* (9). The criterion for polarization is that the growth rate of the first spherical harmonic is larger than 10^−4^ s^−1^, i.e. if Re σ_1_ > 10^−4^ s^−1^. A parameter set is classified as not polarizing if the growth rates of the first mode is negative (which implies that all higher harmonics also decay because the instability is always a long wavelength instability). Note that since we do not distinguish between different GAPs in the model, the *BEM3* knockout amounts to a reduction of the total GAP copy number by the Bem3 copy number: 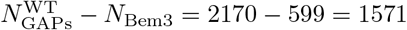. Condition 3 ensures that cell growth alone does not rescue spontaneous polarization of *bem1*Δ. cells (the protein copy numbers are scaled up proportionally to the cell radius).

| # | Polarizes | Experimental mutant / condition | Modified parameters |
| --- | --- | --- | --- |
| **1** | yes | WT | — |
| **2** | no | *bem1*Δ | $N_{\text{Bem1}} = 0$ |
| **3** | no | *bem1*Δ, cell grown to radius 7 $\mu\text{m}$ | $N_{\text{Bem1}} = 0, R = 7 \mu\text{m}$ |
| **4** | yes | *bem1*Δ *bem3*Δ | $N_{\text{Bem1}} = 0,$<br>$N_{\text{GAPs}} = N_{\text{GAPs}}^{\text{WT}} - N_{\text{Bem3}}$ |
| **5** | yes | *bem1*Δ *GAL*-*Cdc42*,<br>in 0.1 % Gal ($\approx 1.5 \times$ Cdc42) | $N_{\text{Bem1}} = 0,$<br>$N_{\text{Cdc42}} = 1.5 \times N_{\text{Cdc42}}^{\text{WT}}$ |
| **6** | yes | *GAL*-*CDC42* after<br>24 h in 0 % Gal (ca. 20% Cdc42<br>remaining) | $N_{\text{Cdc42}} = 0.2 \times N_{\text{Cdc42}}^{\text{WT}}$ |
| 7 | yes | WT (Filtering under the condition<br>that the first mode grows fastest<br>which ensures formation of a single<br>Cdc42-cluster (Klnder et al.,<br>2013), simetimes referred to as<br>“singularity” (Howell et al., 2009).) | — |
| 8 | yes | Cdc42-rit<sup>C</sup> | $k_d = 0$ |
| 9 | (yes) | Cdc42-psy1<sup>TM</sup> (This mutant has<br>been found to polarize in the fission<br>yeast *S. pombe*, but was not yet<br>studied in budding yeast; see text<br>for details) | $k_d = 0,$<br>$D_{m_d} = D_{m_t} = D_{m_{tg}} =$<br>$D_{m_b} = D_{m_{bf}} =$<br>$0.0025 \mu\text{m}^2 \text{s}^{-1}$ |

The growth rate of the first mode serves as an indicator whether the parameter set exhibits spontaneous polarization under the respective conditions. This is then compared to the experimental observation (cells polarize or not). To provide some intuition how much the different experimental findings in different conditions constrain the parameters, we perform the filtering in subsequent steps for groups of conditions (separated by horizontal lines in Table S4).

### Step I: “Core” conditions

We start by filtering for parameter sets consistent with our experiments on WT, *bem1*Δ., and *bem1*Δ. *bem3*Δ. cells with galactose-induced Cdc42 (conditions 1–6 in Table S4). Out of 5 10^6^ randomly generated parameter sets, 7730 fulfill conditions 1–6, corresponding to a fraction of 0.15%. While this may seem like a small fraction, one should keep in mind that the sampled parameter space is 13-dimensional. If the parameters were all equally important and independent, each of them could be varied across 60% of its full range (four orders of magnitude), since 0.15 × 10^−2^ ≈ 0.6^1^3. This illustrates the “curse of dimensionality.” Of course in the real system, some of the parameters are not independent, and some of them are more important (stronger constrained) than others.

### Step II: Growth of a single Cdc42-cluster in WT

Under WT conditions, the first mode (first spherical harmonic) should grow fastest, ensuring formation of a single Cdc42-cluster (“singularity”). Of the 7730 parameter sets obtained in Step I, 1422 fulfill this condition. Note that there might be specific mechanisms to ensure “singularity” in the real system that are not captured by our model. Candidates for this are vesicle-based transport (Freisinger et al., 2013), which we modeled in an effective, coarse-grained way (see Sec. 1.2), and negative feedback via the phosphorylation of Cdc24 by Cla4 (Kuo et al., 2014).

### Step III: Permanently membrane-bound Cdc42

Experiments using a permanently membrane-bound Cdc42 mutant (Cdc42-rit^C^) showed that recycling of Cdc42 via the cytosol is not required for Cdc42 polarization (Bendezú et al., 2015; Kang et al., 2018). We wondered whether our mathematical model could reproduce this experimental finding. The permanent membrane-binding of the Cdc42-rit^C^ mutants is accounted for by setting the Cdc42-detachment rate *k_d_* to zero (see constraint 8 in Table S4). Out of the previously filtered 1422 parameter sets, 358 fulfill this additional condition.

To visualize the filtered parameter sets, we show scatter plots for each pair of parameters (see Fig. S3). This shows how much each parameter is constrained by the experiments and visually highlights pair-wise correlations in the parameter sets.

In summary, we find that parameter sets consistent with the experimental observations cover a large range of parameters. Most parameters can vary over the full four orders of magnitude we sampled. Even the most strongly confined parameters, *k_d_* (Cdc42-GDP membrane dissociation rate) and *k_gt_* (hydrolysis rate of Cdc42 in Cdc42-GAP complexes) each cover more than two orders of magnitude.

**Figure S3.**
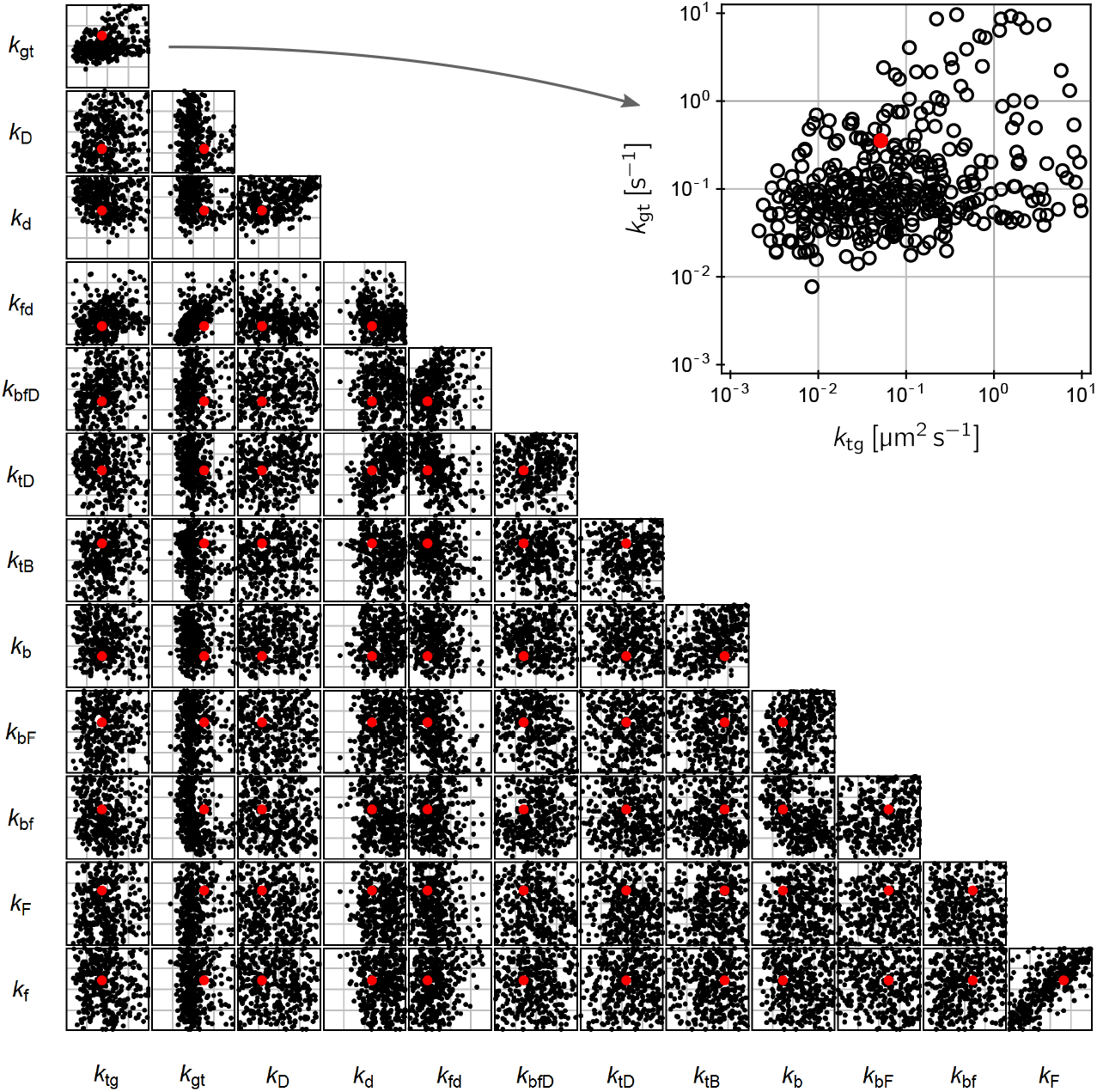
Scatter plots for all pairwise combinations of reaction rates showing the parameter sets that fulfill conditions 1–8 in Table S4 obtained by filtering 5 × 10^6^ randomly generated parameter sets. This parameter set is used in Figs. 2 and 5 and in the Videos 1–6. The red dot marks the parameter set that is closest to the mean of these parameter sets. The inset in the top right corner shows a magnification of the (*k_tg_*, *k_gt_*)-parameter plane. Notably, even the parameter that is constrained the strongest (*k_gt_*, the dissociation rate of Cdc42-GAP complexes) ranges over several order of magnitude.

**Table S5.**
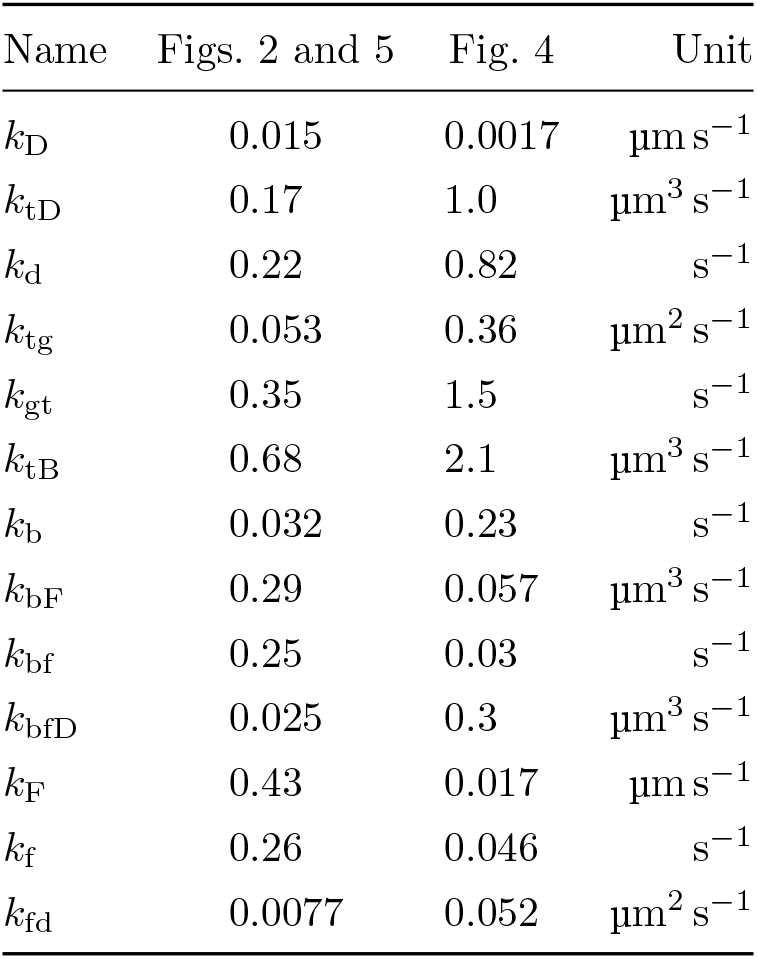
Parameters (kinetic rates described in Tab. S2) obtained by filtering for parameter sets that reproduce the experimental findings. The parameter set used for Figs. 2 and 5 is marked by a red dot in Fig. S3. It was obtained by filtering with conditions 1–8 (Table S4) that correspond to experiments performed with budding yeast and this study and (Bendezú et al., 2015). The parameter set used for Fig. 4, marked by a red dot in Fig. S4, was generated separately to demonstrate that *theoretically* all three mechanisms of polarization encoded in the Cdc42-interaction network can coexist in the same parameter regime. To this end, filtering was performed with conditions 1–6 and 9. The experiment with the membrane-bound, immobile Cdc42 mutant Cdc42-psy1™, corresponding to condition 9, was so far only performed in fission yeast, not in budding yeast.

**Figure S4.**
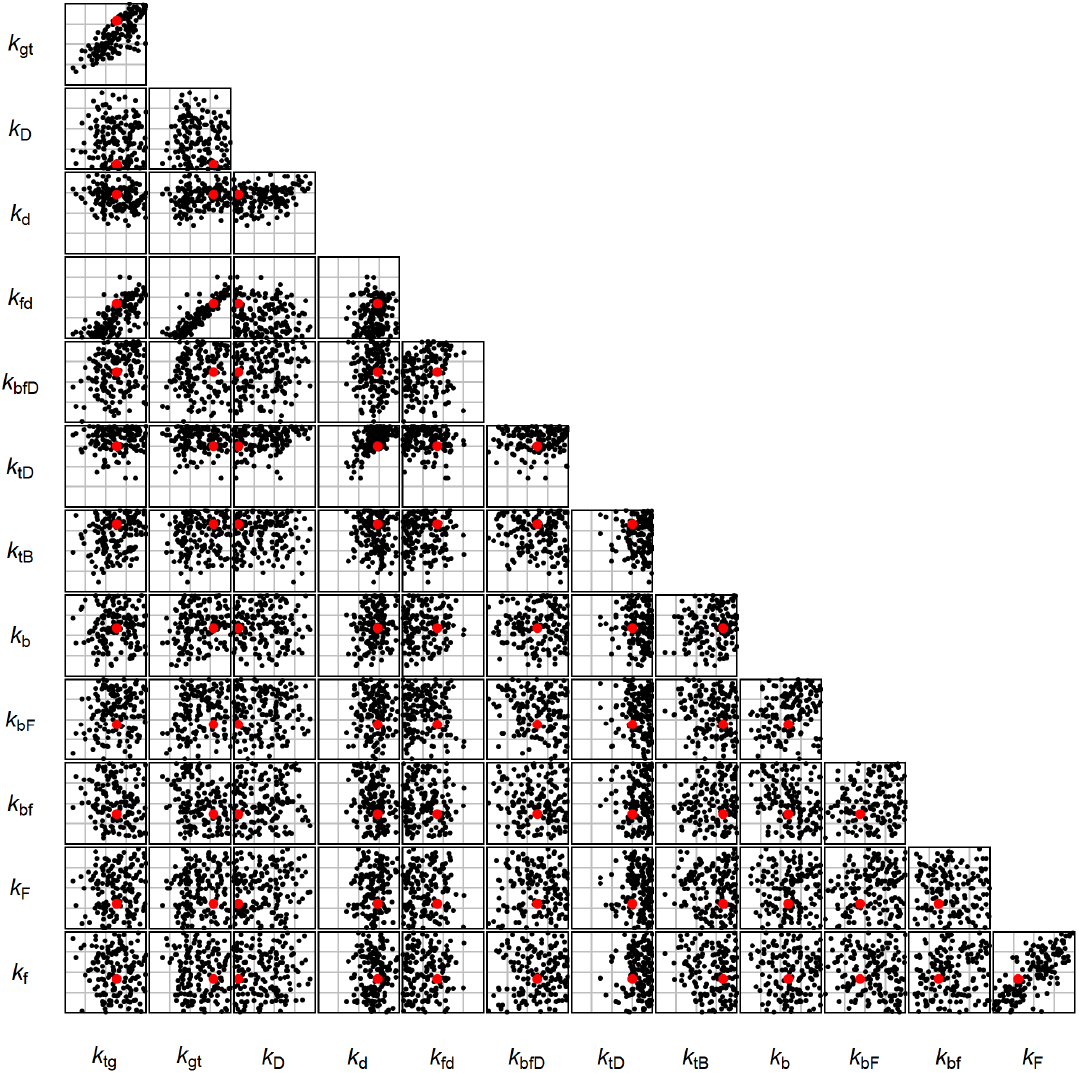
Pairwise scatter plots of the parameter sets that fulfill conditions 1–6 and 9 in Table S4 (out of 5 × 10^6^ randomly generated parameter sets). The red dot marks the parameter set that is closest to the mean. This parameter set is used in Fig. 4.

### Polarization independent of spatial Cdc42 redistribution

The analysis of the Cdc42-polarization machinery in terms of functional subunits carried out in the main text revealed that there might be a polarization mechanism that is independent of spatial Cdc42 redistribution (Fig. 4H). Instead, polarization relies on redistribution of Bem1-GEF complexes and GAP saturation to the polar zone. Cdc42 merely switches between its active and inactive state, while its *total density* remains spatially uniform. To test where this mechanism is operational, we introduce a new condition (9 in Table S4). Under this condition, all Cdc42 is membrane bound (no detachment, *k_d_* = 0) and transport of Cdc42 on the membrane is suppressed by setting the diffusion constants of both active and inactive membrane-bound Cdc42 to same small value (0.0025 μm^2^ s^−1^, based on an estimate from experiments (Valdez-Taubas and Pelham, 2003)). Since for equal diffusion constants the dynamics of Cdc42’s *total density* decouples and is purely diffusive, it remains uniform at all times.

Interestingly, out of the 358 parameter sets obtained in Step III above (fulfilling conditions 1–8 in Table S4 only 33 exhibit spontaneous polarization with immobile Cdc42 (condition 9). This suggests that is unlikely that Cdc42-psy1™ mutants are able to polarize in budding yeast. However, polarization of Cdc42-psy1™ has been found in fission yeast (Bendezú et al., 2015), which exhibits a bi-polar polarization pattern in WT cells. We therefore sampled parameters using only conditions 1–6 and 9, i.e. without the constraint that a single polar cap emerges under WT conditions. The 155 parameter sets found this way are shown in Fig. S4. This demonstrates that the Cdc42-transport independent mechanism could, in principle, be operational in the same parameter regime as the WT and rescue mechanisms. For the phase diagrams shown in Fig. 4 in the main text, we used the parameter set that is closest to the mean of all parameter sets fulfilling conditions 1–6 and 9.

## 6 Supplementary Discussion

### 6.1 Previous experiments explained by our model

#### Permanently membrane-bound Cdc42 mutants

Our model reproduces experiments with Cdc42-rit^C^ fusion chimeras that are permanently membrane-bound (Bendezú et al., 2015), as we have shown in Sec. 5. The only means of Cdc42 transport in these mutants is owing to the slower diffusion of active Cdc42 compared to inactive Cdc42 on the membrane. The formation of complexes between Cdc42-GTP and Gic1/Gic2 has been suggested as a molecular mechanism underlying the slow diffusion of Cdc42-GTP (Kang et al., 2018; Daniels et al., 2018). Indeed, Kang et al. find synthetic lethality of *gic1Δ. gic2Δ*., Cdc42-rit^C^ triple mutants and conclude: *“Therefore, polarization of Cdc42-ritC is likely to be mediated by Gics during the first phase of G1. […] A possible explanation for the synthetic lethality of cdc42-rit^*C*^ gic16. gic26. is that even a slight increase of its mobility (in the absence of Gic1/2) could be more detrimental to polarization of Cdc42-rit^*C*^, which presumably occurs via lateral diffusion and/or GDI-independent exchanges between membrane and cytosol.*” Our model and theoretical analysis suggest that Cdc42-ritC polarization requires differential diffusion of Cdc42-GTP vs Cdc42-GDP on the membrane to achieve redistribution of total Cdc42. The Gics provide this differential diffusion as they effectively reduce the diffusivity of Cdc42-GTP compared to Cdc42-GDP. This corroborates the above explanation by Kang et al. why one of the Gics is required for polarization of cdc42-rit^C^ mutants.

Another strongly membrane-bound Cdc42 mutant, fused to a trans-membrane domain (Cdc42-psy1™), was studied in *S. pombe* (fission yeast) (Bement et al., 2015). Whereas Cdc42-rit^C^ binds to the membrane by an amphipathic helix and therefore can diffuse rather freely on the membrane surface, the transmembrane domain of Cdc42-psy1™ renders this fusion chimera effectively immobile. Strikingly, it was found that Cdc42-psy1™ exhibits polarization of Cdc42 activity without Cdc42 accumulation (Bendezú et al., 2015). Since the Cdc42-polarization machinery shares the key components with budding yeast (see (Chiou et al., 2017) for a review), we hypothesize that this puzzling observation could be explained by our mathematical model of the Cdc42-polarization machinery. In particular, Scd1 and Scd2 (paralogs of Bem1 and Cdc24) facilitate the same scaffold-mediated feedback loop that operates in WT budding yeast. Our mathematical analysis shows that the scaffold-mediated feedback loop in conjunction with GAP-saturation can facilitate polarization independently of Cdc42-redistribution (see Fig. 4H in the main text). Moreover, our parameter study in Sec. 5 above shows that this mechanism can, in principle, be operational in the same regime as the WT mechanism. In contrast to the WT mechanism, our model predicts that polarization with immobile Cdc42 is sensitive to the GAP-Cdc42 copy number ratio because it relies on GAP-saturation in the polar zone; see Fig. 4E,H in the main text.

#### Optogenetic GEF recruitment

In a recent set of experiments in budding yeast, optogenetics was used to transiently recruit Cdc24 to a small membrane patch (Witte et al., 2017). Interestingly it was found that in *bem1*Δ. cells, this optogenetic GEF-localization can induce stable patterns that are maintained even after the optogenetic “stimulus” is removed. This suggests that the *bem1*Δ. cells are in a *subcritical* regime, where a sufficiently strong local perturbation (stimulus) can induce self-sustained polarization. Previous theoretical work has shown that subcriticality is generic in mass-conserving reaction–diffusion systems and that the regimes of stimulus-induced pattern formation are always adjacent to regimes of spontaneous pattern formation. (Trong et al., 2014; Brauns et al., 2018). We therefore expect that the rescue mechanism supports stimulus-induced pattern formation above the critical GAP:Cdc42 threshold in *bem1*Δ cells. Indeed, and in agreement with the experimental findings reported in Witte et al. (2017), numerical simulations of *bem1*Δ. mutants emulating the transient optogenetic recruitment of GEF to a small membrane patch, show that polarization is maintained after the stimulus removed (see Video 5). In conclusion, the rescue mechanism can explain the experimentally found, stimulus-induced polarization of Cdc42, independently of a feedback loop operating on the GEF.

#### Globally enhanced GEF activity rescues Bem1-deletes

Previous experiments have shown that Bem1-mutants that lack the Cdc42-GTP interaction domain but still interact with Cdc24 rescue polarization of *bem1*Δ. cells (Smith et al., 2013; Grinhagens et al., 2020). It has been hypothesized that this is because these Bem1-mutants relieve Cdc42’s autoinhibition and thereby globally increase GEF activity. In accordance with this, the Bem1-independent rescue mechanism predicted by our model can be activated by a global increase of GEF activity (see Fig. 2B in the main text).

In another set of experiments, Bem1’s ability to localize Cdc24 to the zone of high Cdc42-GTP concentration was inhibited by fusing Bem1 to the strongly membrane bound (Woods et al., 2015). Strikingly, this Bem1-Snc2^V39A,M42A^ fusion chimera does not rescue cell polarization. As a reason for this, we hypothesize that the interaction between Bem1-Snc2^V39A,M42A^ and Cdc42 sequesters part of the Cdc42 in the cell and lowers the available amount of Cdc42 below the threshold where the Bem1-independent mechanism is operational. In agreement with this hypothesis, recent results show that sequestration of a fraction of Cdc42 to the membrane by overexpression of the Ccd42-binding protein Gic2 is lethal for *bem1*Δ. bem36. cells (Grinhagens et al., 2020).

### 6.2 Hypothetical feedback via Rsr1 and secondary GEFs

Several proteins from the bud-site selection pathway have been hypothesized to mediate positive feedback loops that may give rise to spontaneous Cdc42-polarization. A hypothetical interaction of the Rsr1-GEF Bud5 with Cdc42 has been suggested recently (Goryachev and Leda, 2017) based on the observation that Bud5 relocalizes from the landmark-determined ring that surrounds the bud-scar to clusters of Cdc42-GTP (Kang, 2001; Marston et al., 2001). This may mediate a feedback loop between Cdc42 and Rsr1 via their respective GEFs Cdc24 and Bud5, that facilitates spontaneous polarization similarly to the Bem1–Cdc42 mutual recruitment mechanism. A similar hypothetical feedback loop mediated by the Rsr1-GEF Bud3 has been suggested (Martin, 2015).

Because proteins suggested to mediate these feedback loops bind to (axial) landmark proteins that localize to the bud scar, it is likely that the feedback would be strongly localized in the vicinity of the bud scar. It was previously observed that Cdc42 polarization in *bem1*Δ.bem36. cells is not preferentially directed to the vicinity of the bud scar. This indicates that landmark associated proteins, such as Rsr1, Bud3, and Bud5 do not play a central role in the rescue mechanism. That these proteins do not play an essential role in the rescue of *bem1*Δ*bem3*Δ cells is further supported by the finding that *bem1Δrsr1Δ* are viable in some strain backgrounds (Smith et al., 2013). Further experiments will be required in the future to elucidate the interplay between the bud-site selection pathway and spontaneous polarization of Cdc42.

## 7 Video Captions

### Video 1

Simulation under WT conditions, see (1) in Table S4. The density of membrane bound Cdc42-GTP concentration, *m*_t_, is plotted on the surface of the sphere that mimics the cell (radius 3.5 μm).

### Video 2

Simulation of a *bem1*Δ*bem3*Δ, see (4) in Table S4. The density of membrane bound Cdc42-GTP concentration, m_t_, is plotted on the surface of the sphere.

### Video 3

Simulation of a *bem1*Δ. mutant with 1.5 Cdc42 overexpression, see (5) in Table S4. The density of membrane bound Cdc42-GTP concentration, *m*_t_, is plotted on the surface of the sphere that mimics the cell (radius 3.5 μm).

### Video 4

Simulation of a wild-type cell with 0.2 Cdc42 underexpression, see (6) in Table S4. The density of membrane bound Cdc42-GTP concentration, m_t_, is plotted on the surface of the sphere that mimics the cell (radius 3.5 μm).

### Video 5

Simulation of Cdc42-rit^C^ mutant, where Cdc42 cannot detach from the membrane, i.e. is transported by lateral diffusion on the membrane alone, see (6) in Table S4. The density of membrane bound Cdc42-GTP concentration, *m_t_*, is plotted on the surface of the sphere that mimics the cell (radius 3.5 μm).

### Video 6

Simulation of a *bem1*Δ. mutant with optognetic GEF recruitment. Left: Visualization of the transient, spatially localized increase in the GEF membrane-attachment rate *k_F_* emulating optogenetic GEF recruitment. The recruitment is switched on in the interval 500 s < *t* < 1000 s and has the shape of a Gaussian pulse with radius 1 μm. Right: Density plot of the membrane bound Cdc42-GTP concentration; cell radius 3.5 μm.

## Footnotes

1 For the four Cdc42 GAPs, a coefficient of variation around 0.14 for cell-to-cell copy-number variability has been reported (Chong et al., 2015). This is on the same order of magnitude as the upper estimate of 25% for the GAP copy number reduction required to activate the Bem1-independent rescue mechanism, suggesting that this mechanism is operational in a fraction of *bem1*Δ cells.

1 To keep the notation compact, we present the detailed derivation for the case of identical cytosolic diffusion constants. Generalization to different cytosolic diffusion constants follows the same steps where each cytosolic component has its own bulk profile *ρ*_*i*_(*r*).

